# In silico design of Phl p 6 variants with altered folding stability significantly impacts antigen processing, immunogenicity and immune polarization

**DOI:** 10.1101/2020.02.26.967265

**Authors:** Petra Winter, Stefan Stubenvoll, Sandra Scheiblhofer, Isabella A Joubert, Lisa Strasser, Carolin Briganser, Soh Wai Tuck, Florian Hofer, Anna Sophia Kamenik, Valentin Dietrich, Sara Michelini, Josef Laimer, Peter Lackner, Jutta Horejs-Hoeck, Martin Tollinger, Klaus R. Liedl, Johann Brandstetter, Christian G. Huber, Weiss Richard

## Abstract

**Introduction:** Protein fold stability has been proposed to represent an intrinsic feature contributing to immunogenicity and immune polarization by influencing the amount of peptide-MHC II complexes (pMHCII). Using *in silico* prediction, we introduced point mutations in proteins that either increase or decrease their fold-stability without altering immunodominant epitopes or changing the overall structure of the protein. Here, we investigated how modulation of the fold-stability of the grass pollen allergen Phl p 6 affects its ability to stimulate immune responses and T cell polarization.

**Methods:** Using the MAESTRO software tool, stabilizing or destabilizing mutations were selected and verified by molecular dynamics simulations. The mutants were expressed in E. coli, purified tag-free, and analyzed for thermal stability and resistance to endolysosomal proteases. The resulting peptides were analysed by degradome assay and mass spectrometry. The structure of the most stable mutant protein was obtained by X-ray crystallography. We evaluated the capacity of the mutants to stimulate T cell proliferation *in vitro*, as well as antibody responses and T cell polarization *in vivo* in an adjuvant-free BALB/c mouse model.

**Results:** Four stabilizing and two destabilizing mutations were identified by MAESTRO. Experimentally determined changes in thermal stability compared to the wild type protein ranged from -5 to +14 °C. Destabilization led to faster proteolytic processing *in vitro*, whereas highly stabilized mutants were degraded very slowly. However, the overall pattern of identified peptides remained very similar. This was confirmed in bone marrow derived dendritic cells that processed and presented the immune dominant epitope from a destabilized mutant more efficiently. *In vivo*, stabilization resulted in a shift in immune polarization as indicated by higher levels of IgG2a and increased secretion of TH1/TH17 cytokines.

**Conclusion:** MAESTRO was very efficient in detecting single point mutations that increase or reduce fold-stability. Thermal stability correlated well with susceptibility to protease resistance and presentation of pMHCII on the surface of dendritic cells *in vitro*. This change in processing kinetics significantly influenced the polarization of T cell responses *in vivo*. Modulating the fold-stability of proteins thus has the potential to optimize and polarize immune responses, which opens the door to more efficient design of molecular vaccines.

## Introduction

An important question for the design and understanding of molecular vaccines is how protein intrinsic factors determine the immunogenic properties of an antigen. Several parameters have been identified that can have a strong impact on the immunogenicity and immune polarizing potential of allergens, like aggregation behavior ^2^, glycosylation ^3^, molecular mimicry ^4^, or enzymatic activity ^5^. However, one of the most important underlying principles is that for a strong immune response, sufficient amounts of peptides have to be presented on major histocompatibility complex molecules to provide optimal stimulation for T cells ^6^. Moreover, the amount of peptide-MHC complexes (pMHC) on the surface of antigen presenting cells (APCs) can influence the polarization of naïve T helper (TH) cells. While in the classic qualitative model, TH cell polarization is mainly determined by cytokines secreted by APCs, there is growing evidence for an important role of strength and duration of the pMHCII - T cell receptor (TCR) interaction. In favor of the quantitative model, it has been consistently shown *in vitro* that low antigen doses promote TH2 polarization, while high antigen doses induce IFN-γ secreting TH1 cells ^7, 8^. More recently, van Panhuys et al. have shown for the first time *in vivo* that the quantity of antigen presented by a dendritic cell (DC) may overrule qualitative signals provided by the same DC, thus shifting T cell polarization from either TH1 to TH2 or vice versa ^9^. Consequently, a model emerges, where depending on the pMHCII – TCR interaction, a DC can induce no response, anergy, TH2 polarization, TH1 polarization, or activation induced cell death with increasing signal strength ^10^. Moreover, it has been shown that TCR signalling strength is also crucial for the induction of Tfh polarization ^11-13^, persistence of Foxp3^+^ Tregs ^14, 15^, and differentiation of TH17 effector cells ^16^. These findings have important implications for the design of novel vaccines, and for our understanding why some proteins are potent TH2 inducers (allergens), while other proteins induce TH1 responses (e.g. viral proteins ^17^).

Based on this concept, the overall fold-stability of an antigen has been suggested as an important protein intrinsic parameter that can influence immunogenicity and immune polarization. Proteins with a high conformational stability are on the one hand more resistant to proteases, which are abundant on skin, mucosal surfaces and in the extracellular matrix ^18, 19^, and on the other hand also frequently display enhanced resistance against proteolytic digestion in the endolysosomal compartment of APCs. Hence, fold stability substantially controls the cell surface density of pMHCII molecules specific for a given antigen, thereby influencing the immune polarization of T cells. Moreover, hyper-stable proteins, which resist proteolysis within the antigen processing compartment, may escape into the cytoplasm of APCs and enter the cross - presentation pathway through the proteasome and finally end up on MHC I. Several studies supporting these ideas have been reviewed by Scheiblhofer et al. ^20^, although their results are somewhat inconsistent due to different methods employed for modulating protein stability and different experimental settings.

There are several methods to manipulate the conformational stability of a protein such as the introduction of cysteine bonds ^21, 22^, pairing of charges ^23^, or chemical cross linking ^24, 25^. However, the outcome of such mutations and the immunological effects of chemical cross linkers are often difficult to predict. In our current work we therefore employed an *in silico* mutagenesis approach using MAESTRO ^26^, a software tool which predicts changes in the free energy (ΔΔG) upon point mutations based on knowledge base potentials and an ensemble of machine learning methods. With this approach, we selected single point mutations that either stabilized or destabilized the grass pollen allergen Phl p 6 without changing known T cell epitopes or the overall structure of the protein. Using these mutant proteins, we investigated the effect of fold-stability on antigen processing and presentation, as well as on immunogenicity and immune polarization *in vitro and in vivo*.

## Materials & Methods

A more detailed version of the Materials and Methods section can be found in the supplement.

### In silico mutagenesis

Stabilizing or destabilizing mutations were selected *in silico* using the MAESTRO algorithm ^26^.

### Expression and purification of recombinant proteins

Wild-type Phl p 6 and its mutants were expressed from pET17b constructs in *Escherichia coli* strain BL21 Star (DE3; Invitrogen/Thermo Fisher Scientific), as described in the supplement.

### Protein characterization and structure determination

Protein folding in solution and thermal denaturation were monitored by circular dichroism (CD), and NMR. The structure was determined by means of x-ray crystallography. Protein flexibility was assessed by using molecular dynamics simulations.

Endotoxin content was determined by *Limulus* amebocyte assay (PYROTELL®-T, Associates of Cape Cod, MA, USA) according to the manufacturer’s instructions. Selected proteins were also tested for masked endotoxin using a NFκB reporter assay based on HEK293 cells overexpressing TLR4, MD-2, and CD14, as previously described ^27, 28^.

### Endolysosomal degradation assay

Endolysosomal degradation assays were performed as previously described ^29^. Briefly, 5µg of protein substrates (Phl p 6 WT, and its variants N16M, E39L, S46Y, and L89G; see results) were mixed with 7.5µg of isolated microsomal fraction from the JAWS II cell line in 50 mmol/L citrate buffer (pH 5.2, or pH 4.5) and 2 mmol/L dithiothreitol. Samples were incubated at 37°C for the indicated time points followed by 5min denaturation at 95°C to stop the reaction. Samples were analyzed on 20% acrylamide gels by SDS-PAGE and coomassie staining followed by densitometric analysis of the full length protein using ImageJ. For Phl p 6 WT, S46Y, and L89G, the pool of generated peptides was additionally assessed by high-performance liquid chromatography-mass spectrometry as described in the supplement.

### Processing of Phl p 6 and Phl p 6 mutants by bone marrow derives dendritic cells and stimulation of Phl p 6–specific T cells

Based on T-cell proliferation data using a Phl p 6 peptide library, peptide 92-106 was chosen for the generation of Phl p 6-specific T cell hybridomas as described in the supplement.

Bone marrow-derived dendritic cells, generated by culturing bone marrow cells in the presence of GM-CSF (GM-CSF BMDCs), were incubated in 96-well U-bottom plates for 16-64h in the presence of 20µg/mL Phl p 6 WT, S46Y, or L89G in GM-CSF containing culture medium (RPMI1640 supplemented with 20 ng/mL GM-CSF, 100 U/mL penicillin, 100 µg/mL streptomycin, 2 mM L-Glutamine, 10% FCS and 50 µM 2-mercaptoethanol), in triplicates. Control wells received medium alone. After washing the cells 2 times with DPBS, 4 x 10^5^ Phl p 6-specific T cell hybridoma cells were added in DMEM, 4mM L-Glu, 10% FCS and co-cultured with protein-loaded BMDCs for another 24h. Standard wells contained BMDCs that were pulsed with different concentrations of peptide 92-106 during the 24h of co-culture. Culture supernatants were removed and analyzed for IL-2 production as a correlate for T cell activation using an ELISA MAX mouse IL-2 set (BioLegend). IL-2 values measured in supernatants from the wells containing BMDCs pulsed with proteins for different time periods were transformed into µM peptide equivalents by interpolating in a standard curve generated from the values obtained from hybridomas co-cultured with BMDCs at indicated peptide concentrations.

### Mice and immunizations

Female 6- to 10-week-old BALB/c mice were obtained from Janvier (Le Genest-Saint-Isle, France) and maintained at the animal facility of the University of Salzburg in a specific pathogen-free environment according to local guidelines. Animal experiments were approved by the Austrian Ministry of Science (permit no. BMWF-66.012/0013-WF/V/3b/2017).

For initial epitope mapping (suppl. Fig. 1) and generation of T cell hybridomas, mice were immunized with 5µg Phl p 6 adsorbed to alum (50% v/v Alu-Gel-S, Serva) by s.c. injection of 200µL divided between two sites on the back. Immunizations were performed on days 0, 19, and 48, and mice were sacrificed one week later.

To assess *in vivo* immunogenicity of the different proteins (Fig. 4) and for repeated epitope mapping (Fig. 5), mice were immunized with 10µg of Phl p 6 or one of the mutant proteins in sterile PBS by means of intradermal injections at two sites on the shaved back (2 x 20µL). Mice were immunized on days 0, 14, and 28. On day 44, blood was taken from retroorbital sinus, mice were sacrificed, splenocytes were prepared and a basophil activation test (BAT) was performed as described in the supplement.

### Antibodies and cytokines

Phl p 6-specific IgG1 and IgG2a were determined by using a luminometric ELISA at serum dilutions of 1:100,000 and 1:100, respectively, lying within the linear range of the assay. Specific serum IgE was assessed by using a β-hexosaminidase release assay. The amount of cell-bound IgE was measured by using a BAT. Both methods are described in detail in the supplement.

Splenocytes from immunized mice were prepared, stained with proliferation dye eFluor 450 (eBioscience/Thermo Fisher Scientific) and cultured in the presence of Phl p 6 or the mutants (20µg/mL) or individual peptides of a 15mer library (GenScript, NJ, USA) with an offset of 3 amino acids (10µg/mL) as described in the supplement. After 4 days, culture supernatants were removed and cells were harvested for flow cytometric analysis as described in the supplement. Cytokine levels in supernatants were analyzed using the LEGENDplex™ Mouse Th Cytokine Panel (13-plex, BioLegend) according to the manufacturer’s instructions.

### Statistics

Statistical difference between groups were analysed by one-way ANOVA followed by Tukey’s post hoc test unless otherwise indicated (Prism 7, GraphPad Software). Data are expressed as means ± SEM. (P-value range is indicated: *P<0.05, **P<0.01, ***P<0.001, and **** P<0.0001).

## Results

### In silico mutagenesis

In an unbiased approach, stabilizing and destabilizing single point mutations were calculated using the MAESTRO stability prediction software ^26^ excluding the dominant T cell epitopes (AA_65-79_ and AA_92-_ 106, see supplementary Fig. 1 and supplementary Table 1) and the candidates were ranked by predicted ΔΔG values (Tables 1 and 2). We found surface exposed as well as buried amino acid exchanges among the top 10 stabilizing mutations, and chose two of each for further analysis (N16M, D52L, E39L, S46Y). On the contrary, the top 10 predicted mutations with destabilizing effect all targeted buried residues, primarily exchanging hydrophobic amino acids within the protein core, potentially leading to complete unfolding as indicated by the high ΔΔG values. Furthermore, the number one candidate mutation was located close to an immunodominant T cell epitope. Therefore, we selected the mutant ranked second (L64G) and performed another scan restricted to surface exposed residues, resulting in more moderate ΔΔG values. As the top two proposed mutations were exchanges to prolines within the alpha helical structure of Phl p 6, we chose mutant L89G (ranked third) for expression and characterization.

**Table 1.** prediction of stabilizing mutations using MAESTRO algorithm. Stabilizing mutations selected for expression and further analysis are marked in red.

| mutant | $\Delta\Delta G_{\text{pred}}$ | c_pred | ASA |
| --- | --- | --- | --- |
| **N16M** | -1,4691 | 0,886 | buried |
| **D52L** | -1,3471 | 0,893 | surface |
| **N16F** | -1,3469 | 0,970 | buried |
| **N16Y** | -1,3052 | 0,982 | buried |
| **E39L** | -1,2817 | 0,890 | surface |
| **S46Y** | -1,2734 | 0,955 | buried |
| **N16L** | -1,2711 | 0,816 | buried |
| **S46F** | -1,2654 | 0,930 | buried |
| **M1R** | -1,2512 | 0,871 | surface |
| **N16W** | -1,2431 | 0,964 | buried |

$\Delta\Delta G_{\text{pred}}$ = predicted change in $\Delta\Delta G$ [kcal/mol]. c\_pred = confidence estimation (0 – 1) of the prediction. Negative values indicate a stabilizing mutation.

**Table 2.** prediction of destabilizing mutations using MAESTRO algorithm. Destabilizing mutations selected for expression and further analysis are marked in green.

| *surface exposed* |  |  | *buried* |  |  |
| --- | --- | --- | --- | --- | --- |
| mutant | $\Delta\Delta G_{\text{pred}}$ | c_pred | mutant | $\Delta\Delta G_{\text{pred}}$ | c_pred |
| L89P | 2,0159 | 0,875 | F91G | 5,1619 | 0,726 |
| A17P | 1,9855 | 0,831 | L64G | 5,0143 | 0,757 |
| L89G | 1,9812 | 0,864 | F19G | 4,7531 | 0,727 |
| A41D | 1,9513 | 0,864 | F87G | 4,6827 | 0,735 |
| D82S | 1,8896 | 0,889 | L60G | 4,6052 | 0,758 |
| A41P | 1,7979 | 0,831 | F91D | 4,5717 | 0,748 |
| D82P | 1,7622 | 0,884 | L64E | 4,5235 | 0,807 |
| A41G | 1,7168 | 0,885 | L11D | 4,4678 | 0,772 |
| A41N | 1,6797 | 0,855 | F38G | 4,4658 | 0,743 |
| A24P | 1,6424 | 0,820 | F42G | 4,4645 | 0,733 |

$\Delta\Delta G_{\text{pred}}$ = predicted change in $\Delta\Delta G$ [kcal/mol]. c\_pred = confidence estimation (0 – 1) of the prediction. Positive values indicate a destabilizing mutation.

### Expression and characterization of Phl p 6 and its variants

Proteins were expressed tag-free in *E. coli* with the exception of mutant L64G, which we were unable to produce, most likely due to its high degree of destabilization, impeding proper folding. Subsequently, proteins were purified by hydrophobic interaction chromatography followed by anion exchange chromatography and finally by size exclusion chromatography. The final purity was determined by SDS-PAGE (suppl. Fig. 2A). After endotoxin removal, all preparations contained less than 1 pg LPS per µg of protein according to *Limulus* amebocyte lysate assay. Proteins used for immunization of mice were additionally tested for masked LPS in a cell-based NF-κB-luciferase reporter gene assay using HEK293 cells overexpressing the LPS receptor subunits TLR4, CD14 and MD-2. Phl p 6 WT and the mutant S46Y were below the detection limit of 0.0025 pg/µg protein; the L89G preparation contained <0.1 pg/µg. Amino acid substitution was confirmed by means of mass spectrometry (suppl. Fig. 2B), and the correct conformation (suppl. Fig. 2C) of the proteins was demonstrated by using CD spectroscopy. Mutant D52L turned out to be truncated and was excluded from further analysis. Stability of the proteins was assessed by CD (Fig. 1A) and NMR (Fig. 1C, WT, L89G, and E39L only). As predicted by MAESTRO, mutants E39L, N16M, and S46Y displayed an increased thermal stability (ΔT 8.8 – 14°C) whereas the destabilized mutant L89G was less stable (ΔT - 5°C), albeit without losing its conformation (Fig. 1B). These predictions, except for N16M, were also confirmed by an accelerated molecular dynamics (aMD) simulation (Fig. 1B and suppl Fig. 3). In the case of N16M the structure was found to be extremely unstable, showing fast unfolding even at the nano-second timescale. To confirm this trend and exclude the possibility of simulation artifacts, the simulations were repeated multiple times with different random starting velocities and also the starting structure was remodeled. Still, the instability of the N16M mutant persisted. NMR confirmed the changes in stability of mutants E39L and L89G, with a visible decrease of signals upon heating in L89G indicating a lower thermal stability compared to the wild-type Phl p 6 For E39L, a high thermal stability could be confirmed, as signal intensities began to diminish at higher temperatures compared to the wild type. (Fig. 1C). A more detailed analysis with x-ray crystallography of mutant S46Y showed that the 3-dimensional structure was remarkably similar to that of the wild-type protein. The superimposition of the crystal structures of S46Y with Phl p 6 WT revealed that the overall fold is identical, with a total root-mean-square deviation of 0.42A. As shown in Fig. 2, the introduction of a large hydrophobic Tyr46 in the α2 helix, pointing into the central hydrophobic cavity contributes to hydrophobic interactions with Phe42 within the α2 helix and also Tyr68 in the neighboring α3 helix). These intra- and inter-helical hydrophobic interactions contribute to the overall structural stabilization.

**Figure 1.**
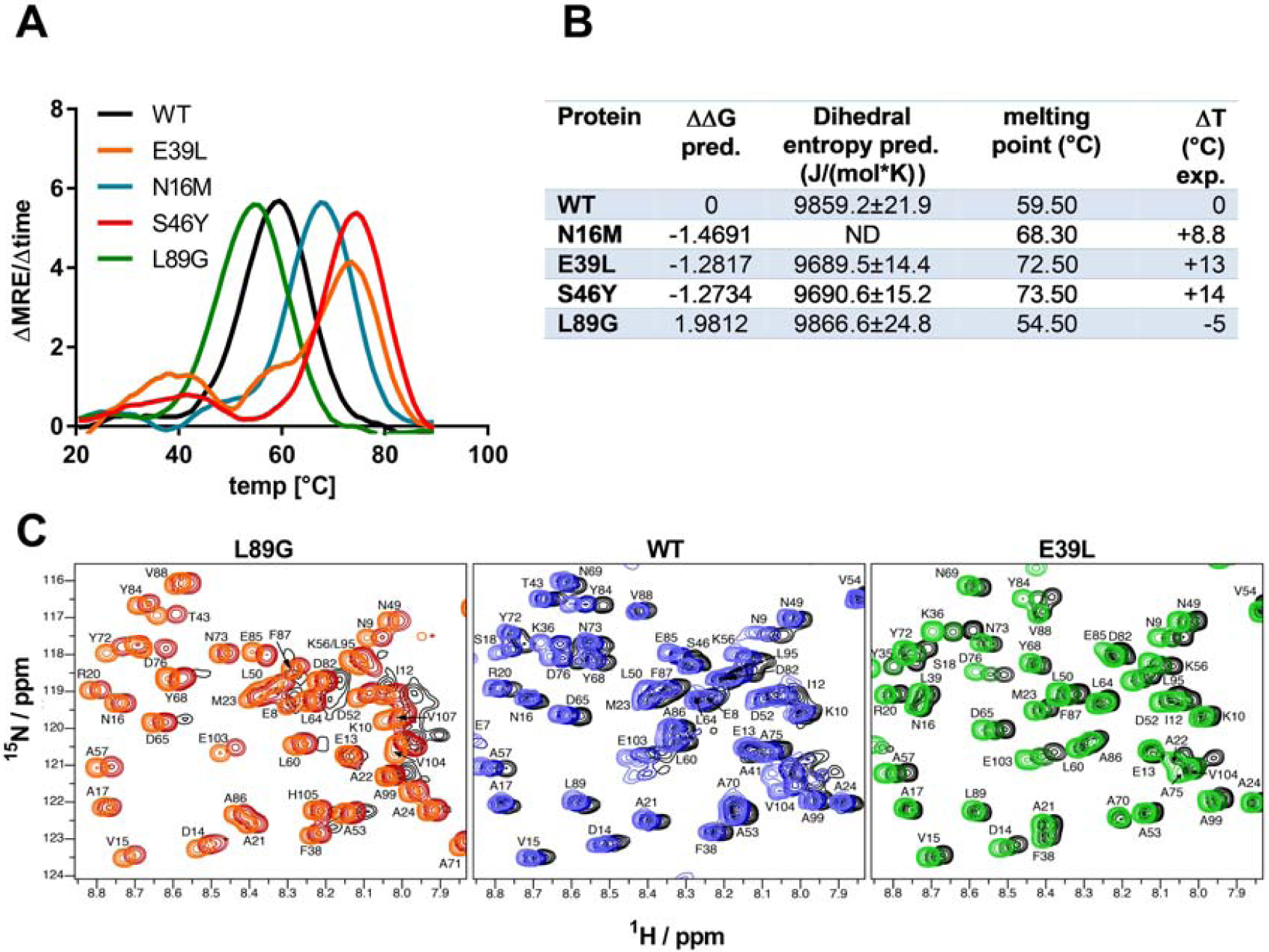
Thermal stability of Phl p 6 wild type (WT) and its mutants. CD spectroscopy was performed at 222nm using a temperature ramp from 20°C to 90°C and the melting point was calculated as the inflection point (peak of the first derivative) of the mean residual ellipticity (MRE) curve (A). Panel B shows the summary of the predicted (pred.) changes in free enthalpy (ΔΔG) and dihedral entropy and the experimentally (exp.) determined thermal melting point and change in melting point (ΔT) compared to the wild type protein. ND = not done. C) Sections of backbone amide ^1^H^15^N HSQC NMR spectra of Phl p 6 recorded at 700MHz at variable temperatures. From left to right: destabilized mutant L89G at 35°C (orange), 40°C (red) and 45°C (black), wild type at 35°C (light blue), 40°C (dark blue) and 45°C (black), and stabilized mutant E39L at 35°C (light green), 40°C (dark green) and 45°C (black). Backbone amide NH cross peaks are labeled.

**Figure 2.**
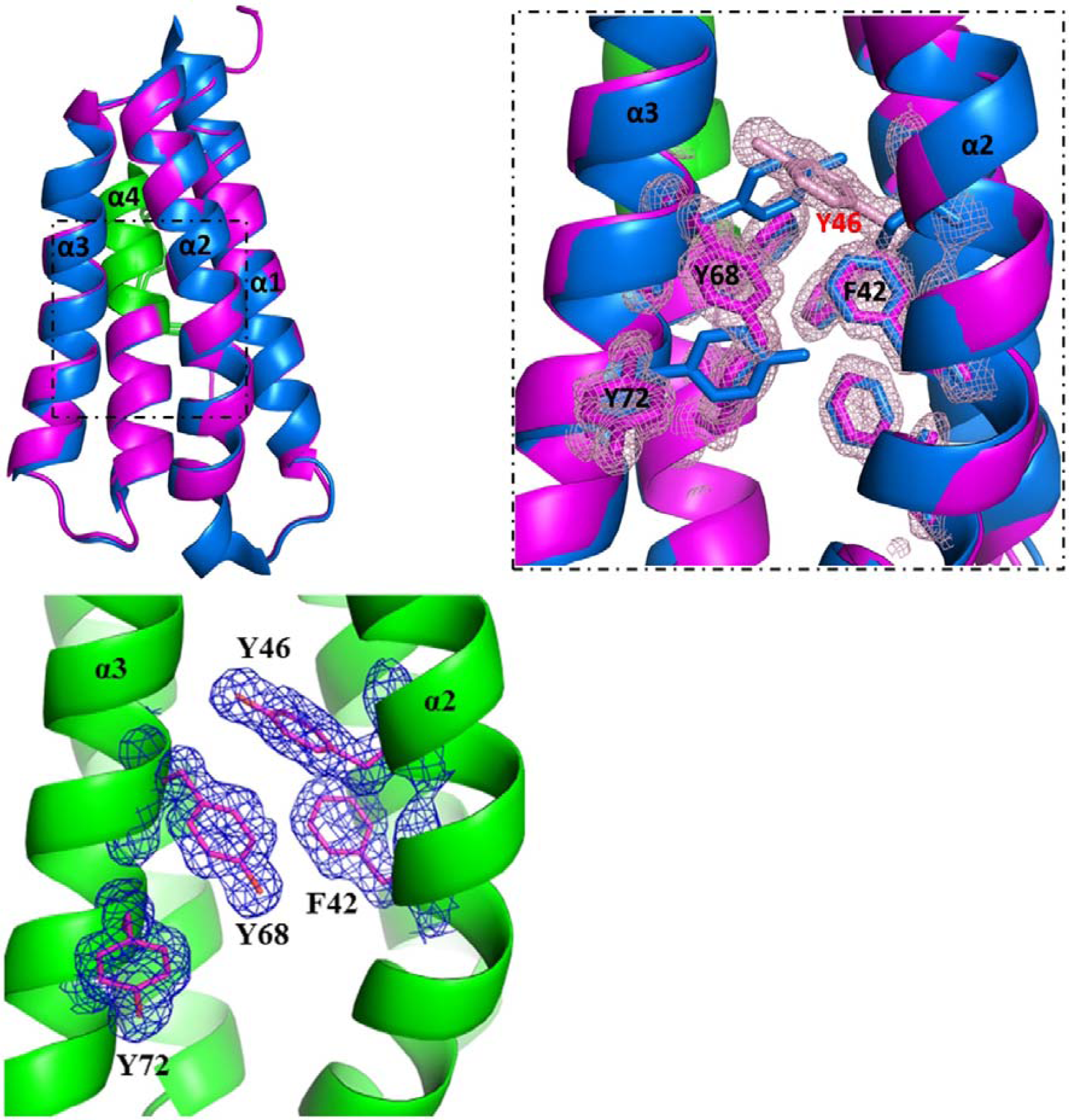
Structural alignment of Phl p 6 wild type (PDB: 1NLX) and mutant S46Y (PDB: 6TRK). The crystal structure of Phl p 6 mutant S46Y was solved at 1.6 Å resolution. Left, mutant S46Y (magenta) has an overall high structural similarity (Cα-RMSD: 0.55 Å) to the wild type protein (blue). The immunodominant epitope (aa91-105) is labelled in green. Right, a close up view of the hydrophobic core. The Tyr46 (pink) was found to complement the aromatic stacking and thereby to improve both the stability of helix α2 harboring Tyr46 as well as the hydrophobic core interaction of the four-helix bundle architecture α1 - α4 of Phl p 6. The density is displayed as a 2Fo-Fc difference map contoured at σ=1.5.

### Fold-stability of Phl p 6 affects proteolytic processing and presentation of antigenic peptides to T cells

To assess how the observed changes in thermal stability affect proteasomal degradation and thus antigen presentation, a degradome assay was performed. The different proteins were incubated with an endosomal extract of the JAWS II dendritic cell line, and degradation of the proteins was determined by SDS-PAGE and Coomassie staining. At a pH of 5.2, the destabilized mutant L89G showed the fastest degradation. Of the stabilized mutants, N16M had no effect on proteolytic stability and resulted in even faster degradation than the WT at some early time-points. In contrast, E39L and S46Y clearly showed delayed proteolytic processing, with S46Y being the most stable protein (Fig 3A, left panel). As expected, at a more acidic pH, proteolysis was more efficient but the picture in general stayed the same (Fig 3A, right panel). As the bulk proteolysis occurred within the first 12h, we repeated the experiment with the WT protein, the destabilized protein L89G and the most stable protein S46Y, and followed the degradation kinetics within the first 12h by quantitating the generated peptides by mass spectrometry. As shown in supplementary Figure 4 (pH 5.2) and 5 (pH 4.5), the general pattern of generated peptides was very similar for the three proteins. However, the processing speed was quite different (Fig 3B). Confirming SDS-PAGE analysis, the N terminus of L89G was processed much faster compared to the WT protein, and also the regions containing the immunodominant epitopes 65-79 and 92-106 were degraded faster at pH 5.2. At pH 4.5, the difference between WT and L89G was less obvious; however, we observed more efficient cutting right next to the N-terminus (position 89-91) of the immunodominant epitope (Fig. 3C). This fits to data from aMD analysis that predicted this region to be more flexible compared to the WT (supplementary Fig. 3) providing an explanation for its higher accessibility to endosomal proteases. We then used T cell hybridomas specific for peptide 92-106 to detect pMHCII complexes on the surface of BMDCs. As shown in Fig. 3D, secretion of IL-2 after stimulation with peptide loaded BMDCs was proportional to the peptide concentration. Using peptide pulsed BMDCs as standard curve, we calculated the pMHCII concentration (as µM peptide equivalents) from BMDCS incubated with the different proteins for 16-64h (Fig. 3E). Confirming the degradome data, peptide 92-106 was much faster and more efficiently processed from the destabilized mutant L89G compared to the WT or S46Y. After 64h, the processing and presentation of peptide 92-106 was 3.3-fold more efficient from WT Phl p 6 and 16.7-fold more efficient from L89G compared to S46Y.

**Figure 3.**
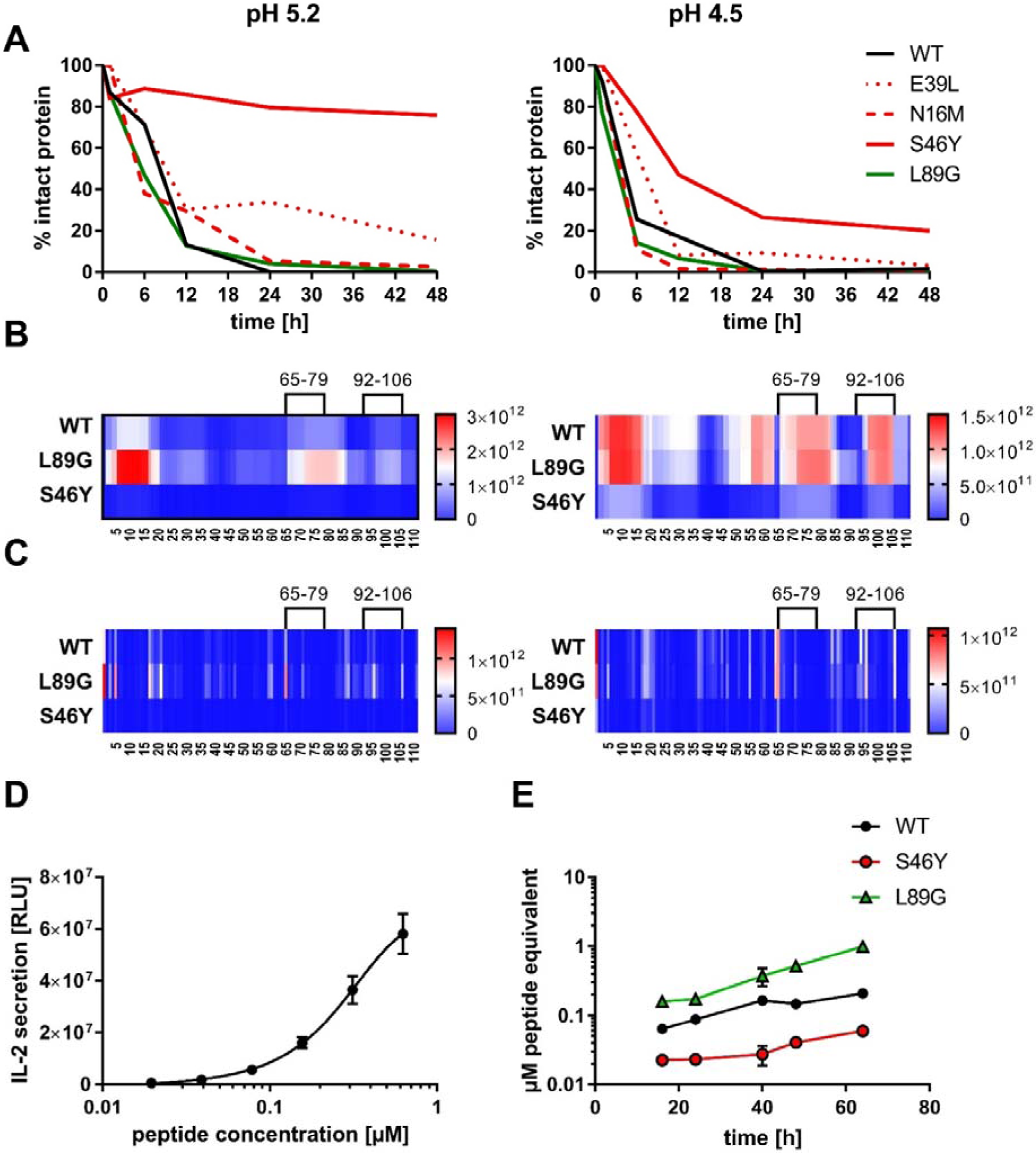
Proteolytic processing and presentation of Phl p 6 and its mutants. A) Proteins were incubated with endolysosomal preparations for 0, 6, 12, 24, and 48h at pH 5.2 (left) and 4.5 (right) and the percentage of remaining intact protein was estimated by SDS-PAGE and Coomassie staining. Peptides generated after 1h, 3h, 6h, and 12h were analyzed by mass spectrometry. Peptides (B) and their respective cutting sites (C) generated earlier during the proteolytic processing are colored in red, whereas peptides that were not detected during the whole experiment are colored in blue. D) Activation of peptide 92-106 specific T cells by BMDCs pulsed with different concentrations of peptide. Data are shown as means±SD (n=3) of relative light units (RLU) of a luminometric ELISA. E) Presentation of peptide 92-106 by BMDCs incubated with 20µg/mL of the different proteins over time.

### Stabilized mutant S46Y shows less IgE binding and induces higher IgG2a compared to the wild type protein and destabilized mutant L89G

To investigate how altered fold stability affects immunogenicity *in vivo*, we immunized BALB/c mice with Phl p 6 WT, stabilized mutant S46Y, and destabilized mutant L89G by intradermal immunization (Fig 4A). To be able to discern modulation of immune polarization induced solely by the stability of the protein, we did not use any adjuvant. Mice immunized with mutant S46Y displayed significantly higher WT-specific IgG2a titers compared to the WT and L89G, which induced the lowest IgG1 as well as IgG2a titers (Fig 4B and C). This effect was not due to major differences in IgG B cell epitopes as sera raised against the mutant proteins recognized the WT molecule with similar efficacy compared to the homologous molecule. As expected, antibodies showed the highest reactivity against the homologous proteins, whereas antibodies raised against the stabilized protein showed the lowest reactivity against the destabilized protein and *vice versa* (Supplementary Figure 5A and B). Similarly, high titered IgE sera raised against the WT protein showed intermediate reactivity with L89G and the lowest reactivity with S46Y (Fig. 4D).

**Figure 4.**
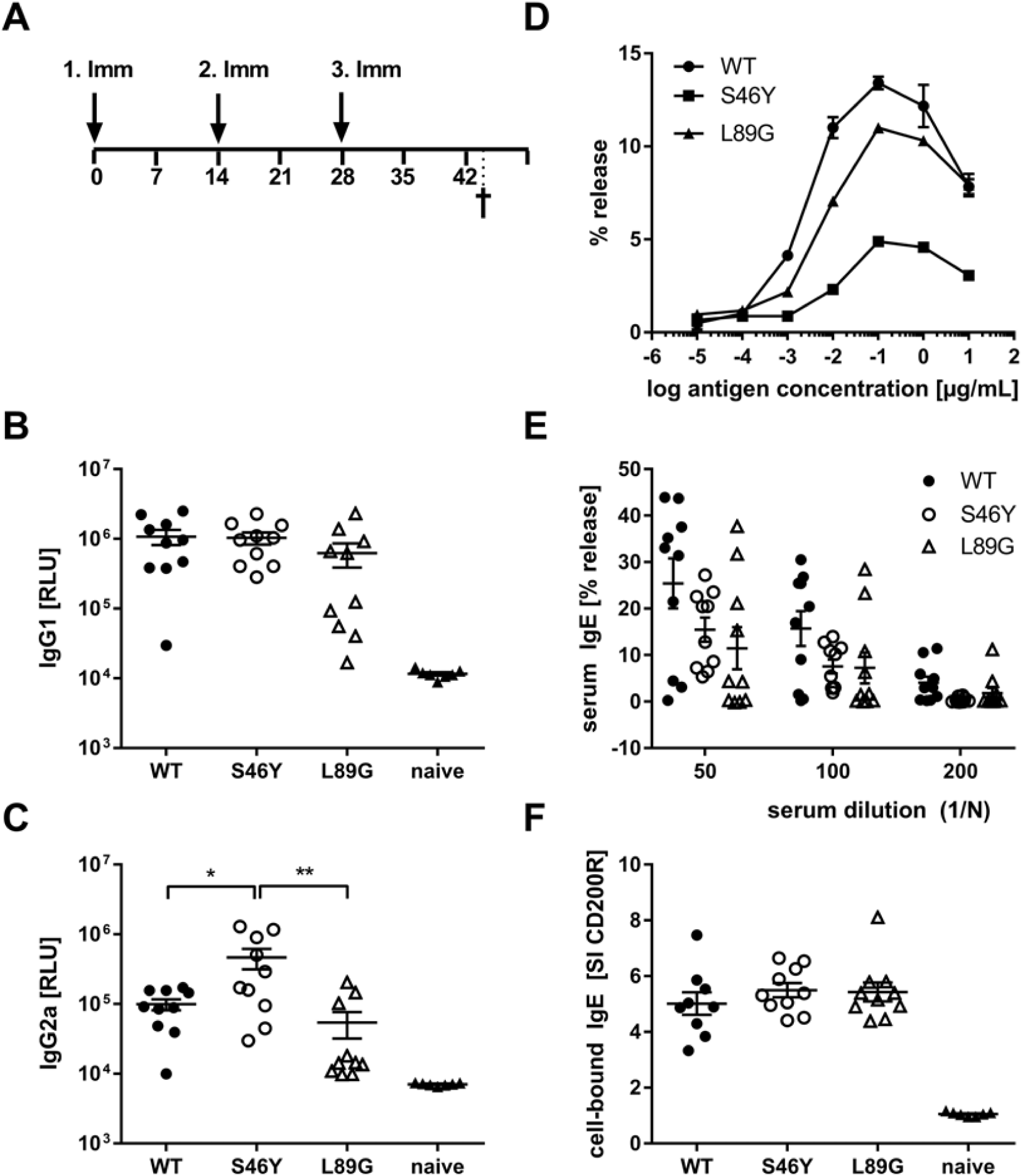
Immunogenicity of Phl p 6 and its mutants. A) Immunization schedule. Phl p 6 specific serum IgG1 (B), IgG2a (C), and IgE (D, E) levels were measured by luminometric ELISA (IgG) and RBL assay (IgE). Samples (n=10) were analyzed at serum dilutions of 1:100000 (IgG1), 1:100 (IgG2a), and as indicated (IgE), lying within the linear range of the assays. D) IgE crosslinking capacity of the different protein at different concentrations using RBL cells loaded with high titered Phl p 6 specific IgE (n=3, means±SD). Data are shown as relative light units (RLU) and % mediator release upon cross-linking with WT protein. E) Cell bound IgE was measured by basophil activation test after *in vitro* stimulation of whole blood samples with 10ng of an equimolar mixture of WT, S46Y, and L89G protein. Data are shown as stimulation indices (SI) of the upregulation of activation marker CD200R.

Although the proteins differed in their IgG2a inducing potential, all groups showed similar levels of serum IgE (RBL assay, Fig. 4E) or cell bound IgE (basophil activation test, Fig. 4F) after *in vitro* restimulation with WT Phl p 6 or an equimolar mixture of WT, S46Y, and L89G, respectively.

It has been shown previously that the fold stability of a protein influences the repertoire of presented peptides and that changing the stability can promote the presentation of hitherto not presented, so-called cryptic epitopes. To confirm our data from the degradome assay that displayed a very similar peptide profile for the different molecules *in vitro*, we again mapped the T cell epitopes of Phl p 6 in mice immunized with the WT protein or the mutants. As shown in Fig. 5, mice immunized with the different Phl p 6 mutants elicited the same profile of T cell responses as the WT molecule. Only the group immunized with the destabilized mutant L89G also displayed a weak response against the region surrounding peptide 13, which may be due to the more relaxed state of L89G (P<0.05 vs S46Y, P=0.057 vs WT).

**Figure 5.**
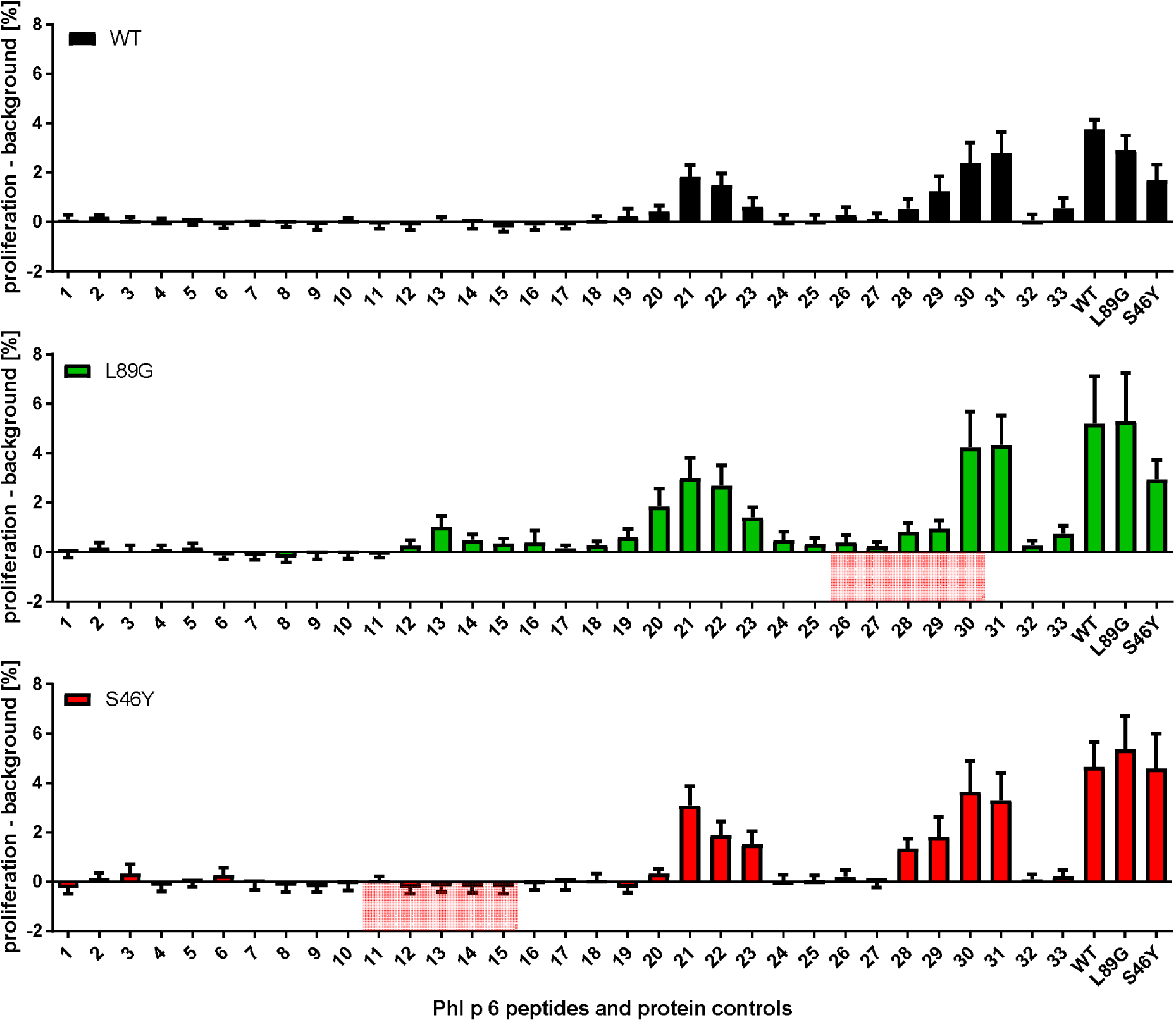
Epitope mapping in mice (n=10) immunized with Phl p 6 WT, and mutants L89G and S46Y. Splenocytes were restimulated with individual 15mer overlapping peptides or the full length protein and proliferation was assessed by flow cytometry. Data are shown as the percentage of proliferating CD4 T cells. Peptides that correspond to the regions containing the respective point mutations in mutants L89G and S46Y are highlighted in red.

### Stabilized mutant S46Y shifts the immune polarization towards inflammatory response types

The amount of pMHCII complexes presented on the surface of APCs has been shown to influence the immune polarization of T cells *in vitro* and *in vivo*. Thus, we hypothesized that the different processing speed of mutant proteins with increased or reduced fold stability might also impact the immune polarization. Therefore, we stimulated splenocytes from immunized mice with the immunodominant peptides 65-79 or 92-106 and measured cytokine secretion in culture supernatants. Although stimulation with peptides 65-79 and 92-106 induced similar T cell proliferation in mice immunized with the wild-type protein and the mutants (Fig. 5), the groups significantly differed in their cytokine profile. In line with the increased IgG2a levels (Fig. 4B), TH1 cytokines TNF-α and IFN-γ were significantly elevated after stimulation with peptide 22 in the group immunized with the stabilized mutant S46Y (Fig. 6A). Similarly, TH17 cytokines IL-17A (not shown), IL-17F, and IL-21 (Fig. 6C) were increased. In contrast, TH2 cytokines IL-4, IL-5, and IL-13 were similar between groups, with the exception of IL-4, which was slightly elevated in the S46Y group after stimulation with peptide 92-106 (Fig. 6B). The regulatory cytokine IL-10 was also elevated in the S46Y group (Fig. 6D), which might be a counter regulation due to the inflammatory response induced by S46Y.

**Figure 6.**
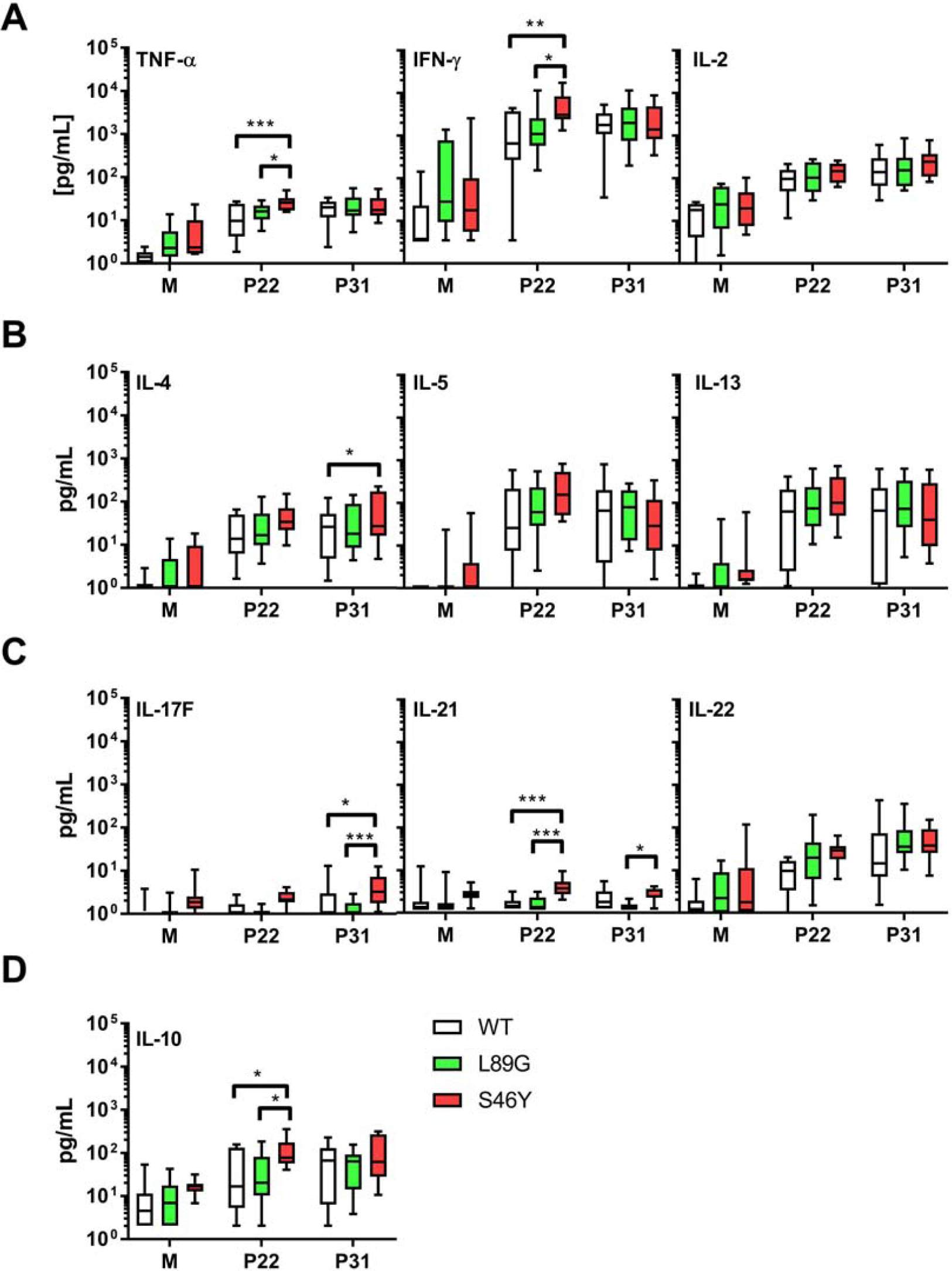
Cytokine levels in supernatants of restimulated splenocytes from mice immunized with Phl p 6 WT, and mutants L89G and S46Y. Cells were restimulated with peptide 65-79 (P22) or 92-106 (P31) or remained unstimulated (medium control, M) for 3 days. Data are shown as box (25^th^ to 75^th^ percentile) and whiskers (min to max) plots (n=10). Statistical differences were assessed by two-way RM ANOVA followed by Tukey’s post hoc test. *** P<0.001, ** P<0.01, * P<0.05.

## Discussion

Enormous efforts have been undertaken to understand and modulate activation and differentiation of Th cells into different effector cell types. The classical qualitative model postulates that MHCII-TCR interaction (signal 1) and co-stimulation via CD28 (signal 2) trigger activation and proliferation of naïve Th cells, whereas cytokines from extrinsic sources such as the APC itself or innate bystander cells largely determine immune polarization. In line with this, individual cytokines or combinations thereof have been identified to drive the development of specific T cell subsets, such as Th1 (IL-12), Th2 (IL-4), Th17 (TGF-β, IL-6, IL-21), Tfh (IL-21), Th9 (TGF-β, IL-4), iTreg (TGF-β, IL-2), and Tr1 (IL-10, IL-27) cells ^30^. However, accumulating evidence suggests that also the quantity and the quality of the TCR signalling induced by pMHCII stimulation plays a crucial role in T cell polarization. In support of this quantitative model, it has been shown that high doses of antigen increase the number of pMHCII complexes presented on the surface of the APC affecting TCR signal strength and/or duration. In summary, these studies suggest that weak TCR:pMHCII interactions favour Th2 or Tfh responses, whereas strong interactions induce Th1 responses (reviewed in ^31^). This model has important implications on our understanding why some proteins, such as allergens, seem to have intrinsic features that drive a certain type of immune polarization. We have previously suggested that protein fold stability represents such an intrinsic feature that has significant impact on the immunogenicity and allergenicity of proteins ^1^. Efficient antigen processing relies on antigen unfolding within the endolysosome. This process is mediated by endolysosomal acidification, and proteolytic cleavage of proteins by resident proteases, which preferentially degrade proteins in an unfolded state, facilitating access to the protein backbone ^32^. However, there is very little *in vivo* data available focusing on the impact of fold stability on polarization of the immune response. Furthermore, studies investigating alterations of immune responses caused by modulated antigen stability have often been contradictory ^20^.

In our current study of Phl p 6, we used an unbiased *in silico* approach to predict stability changes upon the introduction of single point mutations leaving known T cell epitopes and the overall structure unchanged. Indeed, the predicted changes in ΔΔG correlated well with the thermal denaturation as measured by CD spectroscopy, as well as aMD simulations and NMR measurements. In the destabilized mutant L89G, a surface exposed leucine was replaced by a glycine, thereby removing the side chain at this position, giving adjacent amino acids additional flexibility, which was confirmed by aMD simulations. In mutant S46Y, a buried polar amino acid was replaced by a hydrophobic amino acid. Such replacements have been referred to as “hydrophobic core packing” and reported to have stabilizing effects on the protein structure by increasing the overall hydrophobicity within the protein core ^23^. Furthermore, the removal of cavities, by introduction of point mutations, has been shown to have stabilizing effects as well ^n33-35^. During antigen processing, protein unfolding represents an indispensable step, as proteases require access to the backbone of the proteins for efficient cleavage ^32^. Following cellular uptake, antigens are exposed to a continuous drop in pH from mild acidic conditions in the early endolysosome down to pH 4.0 in terminal lysosomes. Fold stability, in particular resistance to pH-mediated denaturation, determines the extent and time-point of unfolding during lysosomal acidification. It has been shown that early endosomes (pH∼6) and terminal lysosomes (pH ∼4,0 - 4,5) are poor in MHC II molecules whereas high quantities of MHC II are found in the late endosome (pH ∼4,5 - 5,5), where the majority of MHC II loading takes place ^36^. We propose a model in which efficient antigen processing requires high fold stability in the early endosome, allowing the accumulation of intact antigen in the late endosome where it becomes sensitive to pH mediated unfolding, facilitating efficient proteolytic degradation ^1^. Thermal stability of Phl p 6 and its mutants correlated remarkably well with the speed of proteolytic digestion using endosomal extracts (Fig. 3A and B) with the exception of N16M, which despite a considerable shift in melting point (+8.8°C) did not display enhanced protease resistance. This is in line with aMD simulations that indicated N16M to be extremely unstable, showing fast unfolding even at the nano-second timescale. Thus, aMD simulations may be a very useful tool to eliminate false positives from the MAESTRO results. To assess the presentation of the immunodominant T cell epitope AA92-106 in the context of proteins with different fold stability, we used BMDCs and a T cell hybridoma line specific for the respective pMHCII complex. IL-2 secretion by the hybridoma cells directly correlated with the amount of peptide loaded on BMDCs and thus represents a sensitive biological surrogate parameter to determine the amount of pMHCII complexes on the surface of APCs. We found that peptide 92-106 was much faster and more efficiently processed from L89G compared to the WT or S46Y, which is in line with enhanced susceptibility to proteases of L89G observed in the degradome assay. Data on the protein dynamics from aMD simulations indicated that the introduction of glycin 89 enhanced the flexibility in the alpha helix (AA91-96) adjacent to the epitope (suppl. Fig. 3) facilitating access to additional cleavage sites (Fig. 3C and D). In contrast, presentation of peptide 92-106 was almost abrogated in the most rigid mutant S46Y. Thus, we could demonstrate that the efficacy of presentation for a given epitope is largely dependent on the fold stability of the protein.

Since we also wanted to investigate the impact of fold stability on the immunogenicity of the generated Phl p 6 derivatives in an adjuvant free setting *in vivo*, it was crucial to get rid of any contaminations, which might modulate the antigen triggered immune response, in particular remaining LPS within the protein solutions. It was previously shown, that LPS concentrations of 0.2 ng/ml are sufficient to activate human monocytes and CD1c^+^ DCs *in vitro* ^27^. LPS from our recombinant proteins was therefore removed to less than 0.1 pg per µg protein, which corresponds to < 2 pg/ml in the *in vitro* experiments and to < 1 pg LPS per mouse for each immunization in the *in vivo* experiments. Therefore, we can exclude LPS induced effects in our experiments. Despite the much slower protein degradation *in vitro*, S46Y turned out to be significantly more immunogenic in terms of IgG2a induction, while IgG1 levels, serum and cell bound IgE remained unaltered (Fig. 4). Although stabilization (and to a lesser extent destabilization) significantly reduced the binding of Phl p 6-specific IgG and IgE antibodies (suppl. Fig. 6), sera from mice immunized with S46Y recognized WT Phl p 6 with slightly lower (IgG1) or very similar (IgG2a) efficacy. Therefore, the changes in Phl p 6-specific antibody levels cannot be explained by changes in conformational B cell epitopes. The observed increase in IgG2a antibodies is in line with a significantly altered cytokine response, showing a shift towards secretion of Th1 and Th17 cytokines after stimulation with the immunodominant peptides (Fig. 6). Various other studies have investigated the effect of fold stability on the immunogenicity of proteins and the subsequent T cell polarization with divergent results. Some reported a higher immunogenicity of stabilized proteins with respect to antibody titers ^37, 38^. Others have found exactly the opposite, namely a reduced immunogenicity of stabilized variants ^39, 40^ and an increased immune response due to an elevated peptide presentation in destabilized proteins ^32^. Further, the immunogenicity is reduced upon protein denaturation ^41^ and increased protease susceptibility ^24^. With regards to T cell polarization, a stabilized form of Bet v 1, including a short Mal d 1 sequence, elicited an increased Th1 response compared to the wild type Bet v 1 ^38^. In contrast, we found an increased Th2 response for Bet v 1 containing stabilizing point mutations in comparison to WT Bet v 1 ^1^. For hen egg lysozyme (HEL), induction of an increased amount of IL-4 could be detected for the destabilized variant, suggesting an increased Th2 response. However, the amount of IFN-γ was also elevated after immunization with the destabilized HEL and not detectable for the stabilized form, indicating an overall higher immune response ^42^. We postulate a model ^20^ where proteins with high protease susceptibility are rapidly degraded in the early endosome, resulting in low numbers of pMHCII complexes to be loaded in the MIIC. Only proteins with sufficient stability survive long enough to reach the MIIC where they are processed and loaded on MHC-II. However, hyperstable proteins that fail to unfold in the MIIC cannot be processed at all and are thus not immunogenic ^1^. Consequently, an optimal stability will generate a high density of pMHCII, which is important for TCR signaling and induction of a Th1 response ^10, 43^. Deviation from this optimum – in both directions – can result in Th2 polarization. This would explain, why in different studies, depending on the initial stability of the used protein, additional stabilization can lead to divergent results.

Mutations in the native protein ^44, 45^, very high antigen concentrations, or unusual protein conformations ^46, 47^ have been shown to induce presentation of so-called cryptic epitopes, that are usually not presented. In this context, we investigated whether changes in fold-stability, which resulted in substantial changes in the processing kinetics *in vitro*, would also change the epitope usage *in vivo*. However, as shown in Fig. 5, the epitope usage between WT, L89G, and S46Y remained largely unchanged, with the exception of the region around peptide 13, which was only recognized by T cells from mice immunized with L89G, indicating that the more relaxed conformation of L89G resulted indeed in the presentation of a cryptic epitope.

In summary, MAESTRO was very efficient in detecting single point mutations that increase or reduce fold-stability, at least for small, alpha-helical proteins such as Bet v 1 ^1^ or Phl p 6 and aMD simulations could be successfully used to eliminate false positives. Thermal stability correlated well with susceptibility to protease resistance and presentation of pMHCII on the surface of BMDCs *in vitro*. Surprisingly, more efficient processing in these *in vitro* systems did not correlate with enhanced immunogenicity and Th1 polarization *in vivo*. On the contrary, the more stable mutant S46Y turned out to be more immunogenic and Th1 polarizing than the destabilized mutant L89G. Potentially, in the *in vivo* setting, either the enhanced protease resistance of S46Y provides the molecule with the advantage of longer extracellular survival (depot effect), or the *in vivo* situation requires slower antigen processing compared to the *in vitro* assays. This applies for intradermal injections where skin dendritic cells take up the antigen and migrate to skin draining lymph nodes during a time-period of 24-48h before they encounter their matching T cells. Additional experiments are necessary to test this hypothesis. Taken together, we have shown that *in silico* evaluation of fold stability modulated proteins has the potential to optimize and polarize immune responses, which opens the door to more efficient design of molecular vaccines.

## Acknowledgements

This work was supported by the Austrian Science Fund (FWF) projects #P26997B13, #P30737, and #W1213 and the Austrian Research Promotion Agency FFG (West Austrian BioNMR 858017).

## Author contributions

PW, S Stubenvoll, S Scheiblhofer, IAJ and RW expressed, purified and characterized recombinant proteins and performed *in vitro* degradation and presentation assays and *in vivo* experiments. LS, CB, and CH performed mass spectrometry and analyzed the respective data, SWT and JB performed X-ray crystallography and analyzed the structures, FH, ASK, and KL developed aMD algorithms and performed the respective analyses. VF and MT expressed proteins for NMR and performed NMR spectroscopy. JL and PL designed the MAESTRO algorithm, modeled the Phl p 6 structure for analysis and performed MAESTRO analyses. SM and JHH performed masked LPS assays and analysed the respective data. S Scheiblhofer and RW designed the study and wrote the manuscript. All authors reviewed, revised and approved the final manuscript.

## References

1. Machado Y, Freier R, Scheiblhofer S, Thalhamer T, Mayr M, Briza P, et al. Fold stability during endolysosomal acidification is a key factor for allergenicity and immunogenicity of the major birch pollen allergen. J Allergy Clin Immunol 2016; 137:1525–34.

2. Rosenberg AS. Effects of protein aggregates: an immunologic perspective. AAPS J 2006; 8:E501–7.

3. Almond RJ, Flanagan BF, Antonopoulos A, Haslam SM, Dell A, Kimber I, et al. Differential immunogenicity and allergenicity of native and recombinant human lactoferrins: role of glycosylation. Eur J Immunol 2013; 43:170–81.

4. Thomas WR. Molecular mimicry as the key to the dominance of the house dust mite allergen Der p 2. Expert Rev Clin Immunol 2009; 5:233–7.

5. Sehgal N, Custovic A, Woodcock A. Potential roles in rhinitis for protease and other enzymatic activities of allergens. Curr Allergy Asthma Rep 2005; 5:221–6.

6. Lever M, Maini PK, van der Merwe PA, Dushek O. Phenotypic models of T cell activation. Nat Rev Immunol 2014; 14:619–29.

7. Hosken NA, Shibuya K, Heath AW, Murphy KM, O’Garra A. The effect of antigen dose on CD4+ T helper cell phenotype development in a T cell receptor-alpha beta-transgenic model. J Exp Med 1995; 182:1579–84.

8. Constant S, Pfeiffer C, Woodard A, Pasqualini T, Bottomly K. Extent of T cell receptor ligation can determine the functional differentiation of naive CD4+ T cells. J Exp Med 1995; 182:1591–6.

9. van Panhuys N, Klauschen F, Germain RN. T-cell-receptor-dependent signal intensity dominantly controls CD4(+) T cell polarization In Vivo. Immunity 2014; 41:63–74.

10. van Panhuys N. TCR Signal Strength Alters T-DC Activation and Interaction Times and Directs the Outcome of Differentiation. Front Immunol 2016; 7:6.

11. Deenick EK, Chan A, Ma CS, Gatto D, Schwartzberg PL, Brink R, et al. Follicular helper T cell differentiation requires continuous antigen presentation that is independent of unique B cell signaling. Immunity 2010; 33:241–53.

12. Fazilleau N, McHeyzer-Williams LJ, Rosen H, McHeyzer-Williams MG. The function of follicular helper T cells is regulated by the strength of T cell antigen receptor binding. Nat Immunol 2009; 10:375–84.

13. Baumjohann D, Preite S, Reboldi A, Ronchi F, Ansel KM, Lanzavecchia A, et al. Persistent antigen and germinal center B cells sustain T follicular helper cell responses and phenotype. Immunity 2013; 38:596–605.

14. Gottschalk RA, Hathorn MM, Beuneu H, Corse E, Dustin ML, Altan-Bonnet G, et al. Distinct influences of peptide-MHC quality and quantity on in vivo T-cell responses. Proc Natl Acad Sci U S A 2012; 109:881–6.

15. Gottschalk RA, Corse E, Allison JP. TCR ligand density and affinity determine peripheral induction of Foxp3 in vivo. J Exp Med 2010; 207:1701–11.

16. Iezzi G, Sonderegger I, Ampenberger F, Schmitz N, Marsland BJ, Kopf M. CD40-CD40L cross-talk integrates strong antigenic signals and microbial stimuli to induce development of IL-17-producing CD4+ T cells. Proc Natl Acad Sci U S A 2009; 106:876–81.

17. Dadaglio G, Moukrim Z, Lo-Man R, Sheshko V, Sebo P, Leclerc C. Induction of a polarized Th1 response by insertion of multiple copies of a viral T-cell epitope into adenylate cyclase of Bordetella pertussis. Infect Immun 2000; 68:3867–72.

18. Turk B. Targeting proteases: successes, failures and future prospects. Nat Rev Drug Discov 2006; 5:785–99.

19. Cudic M, Fields GB. Extracellular proteases as targets for drug development. Curr Protein Pept Sci 2009; 10:297–307.

20. Scheiblhofer S, Laimer J, Machado Y, Weiss R, Thalhamer J. Influence of protein fold stability on immunogenicity and its implications for vaccine design. Expert Rev Vaccines 2017; 16:479–89.

21. Mansfeld J, Vriend G, Dijkstra BW, Veltman OR, Van den Burg B, Venema G, et al. Extreme stabilization of a thermolysin-like protease by an engineered disulfide bond. J Biol Chem 1997; 272:11152–6.

22. Pantoliano MW, Ladner RC, Bryan PN, Rollence ML, Wood JF, Poulos TL. Protein engineering of subtilisin BPN’: enhanced stabilization through the introduction of two cysteines to form a disulfide bond. Biochemistry 1987; 26:2077–82.

23. Lee B, Vasmatzis G. Stabilization of protein structures. Curr Opin Biotechnol 1997; 8:423–8.

24. Delamarre L, Couture R, Mellman I, Trombetta ES. Enhancing immunogenicity by limiting susceptibility to lysosomal proteolysis. J Exp Med 2006; 203:2049–55.

25. Arancibia S, Del Campo M, Nova E, Salazar F, Becker MI. Enhanced structural stability of Concholepas hemocyanin increases its immunogenicity and maintains its non-specific immunostimulatory effects. Eur J Immunol 2012; 42:688–99.

26. Laimer J, Hofer H, Fritz M, Wegenkittl S, Lackner P. MAESTRO--multi agent stability prediction upon point mutations. BMC Bioinformatics 2015; 16:116.

27. Schwarz H, Schmittner M, Duschl A, Horejs-Hoeck J. Residual endotoxin contaminations in recombinant proteins are sufficient to activate human CD1c+ dendritic cells. PLoS One 2014; 9:e113840.

28. Schwarz H, Gornicec J, Neuper T, Parigiani MA, Wallner M, Duschl A, et al. Biological Activity of Masked Endotoxin. Sci Rep 2017; 7:44750.

29. Egger M, Jurets A, Wallner M, Briza P, Ruzek S, Hainzl S, et al. Assessing protein immunogenicity with a dendritic cell line-derived endolysosomal degradome. PLoS One 2011; 6:e17278.

30. Martinez-Sanchez ME, Huerta L, Alvarez-Buylla ER, Villarreal Lujan C. Role of Cytokine Combinations on CD4+ T Cell Differentiation, Partial Polarization, and Plasticity: Continuous Network Modeling Approach. Front Physiol 2018; 9:877.

31. Tubo NJ, Jenkins MK. TCR signal quantity and quality in CD4(+) T cell differentiation. Trends Immunol 2014; 35:591–6.

32. Thai R, Moine G, Desmadril M, Servent D, Tarride JL, Menez A, et al. Antigen stability controls antigen presentation. J Biol Chem 2004; 279:50257–66.

33. Eriksson AE, Baase WA, Zhang XJ, Heinz DW, Blaber M, Baldwin EP, et al. Response of a protein structure to cavity-creating mutations and its relation to the hydrophobic effect. Science 1992; 255:178–83.

34. Lee B. Estimation of the maximum change in stability of globular proteins upon mutation of a hydrophobic residue to another of smaller size. Protein Sci 1993; 2:733–8.

35. Rashin AA. Aspects of protein energetics and dynamics. Prog Biophys Mol Biol 1993; 60:73–200.

36. van Niel G, Wubbolts R, Stoorvogel W. Endosomal sorting of MHC class II determines antigen presentation by dendritic cells. Curr Opin Cell Biol 2008; 20:437–44.

37. Sojikul P, Buehner N, Mason HS. A plant signal peptide-hepatitis B surface antigen fusion protein with enhanced stability and immunogenicity expressed in plant cells. Proc Natl Acad Sci U S A 2003; 100:2209–14.

38. Wallner M, Hauser M, Himly M, Zaborsky N, Mutschlechner S, Harrer A, et al. Reshaping the Bet v 1 fold modulates T(H) polarization. J Allergy Clin Immunol 2011; 127:1571–8 e9.

39. Ohkuri T, Nagatomo S, Oda K, So T, Imoto T, Ueda T. A protein’s conformational stability is an immunologically dominant factor: evidence that free-energy barriers for protein unfolding limit the immunogenicity of foreign proteins. J Immunol 2010; 185:4199–205.

40. So T, Ito HO, Koga T, Watanabe S, Ueda T, Imoto T. Depression of T-cell epitope generation by stabilizing hen lysozyme. J Biol Chem 1997; 272:32136–40.

41. Mine Y, Zhang JW. Comparative studies on antigenicity and allergenicity of native and denatured egg white proteins. J Agric Food Chem 2002; 50:2679–83.

42. So T, Ito H, Hirata M, Ueda T, Imoto T. Contribution of conformational stability of hen lysozyme to induction of type 2 T-helper immune responses. Immunology 2001; 104:259–68.

43. Corse E, Gottschalk RA, Allison JP. Strength of TCR-peptide/MHC interactions and in vivo T cell responses. J Immunol 2011; 186:5039–45.

44. Goldsteins G, Persson H, Andersson K, Olofsson A, Dacklin I, Edvinsson A, et al. Exposure of cryptic epitopes on transthyretin only in amyloid and in amyloidogenic mutants. Proc Natl Acad Sci U S A 1999; 96:3108–13.

45. Topalian SL, Gonzales MI, Ward Y, Wang X, Wang RF. Revelation of a cryptic major histocompatibility complex class II-restricted tumor epitope in a novel RNA-processing enzyme. Cancer Res 2002; 62:5505–9.

46. Lanzavecchia A. How can cryptic epitopes trigger autoimmunity? J Exp Med 1995; 181:1945–8.

47. Warnock MG, Goodacre JA. Cryptic T-cell epitopes and their role in the pathogenesis of autoimmune diseases. Br J Rheumatol 1997; 36:1144–50.

