## Supplementary Data for "In silico design of Phl p 6 variants with altered folding stability significantly impacts antigen processing, immunogenicity and immune polarization"

### Supplementary methods:

#### In silico mutagenesis

Stabilizing or destabilizing mutations were selected *in silico* using the MAESTRO algorithm ^1^ for predicting the influence of single point mutations or combinations thereof on protein stability. The change in stability predictions are based on the PDB entry 1NLX chain L. The structurally unresovled residues at the N- and C-terminus were modeled using the UCSF Chimera ^2^ interface to MODELLER ^3^. The final model was subjected to a MAESTRO greedy search, excluding the known major T cell epitopes 65-79 (DEVYNAAYNAADHAA) and 92-106 (SEALRIIAGTPEVHA). Mutations with a relative accessible surface area or <30% were defined as buried residues and >30% as surface exposed residues.

#### Expression and purification of recombinant proteins

Wild-type Phl p 6 and its mutants were expressed from pET17b constructs in *Escherichia coli* strain BL21 Star (DE3) (Invitrogen/Thermo Fisher Scientific). Cells were grown in auto-inducing ZYM 5052 medium ^4^ supplemented with 100mg/L ampicillin in baffled flasks at 37°C for 20h. Cells were lysed in 6mL lysis buffer (25mM imidazole, 0.5mM EDTA, pH 7.4) per gram pellet weight and 300µg/mL lysozyme was added. After stirring for 30min at RT, bacterial cells were lysed by three freeze/thaw cycles at 70°C. DNA was digested by addition of 100µg DNAse I (Roche) per gram pellet weight and stirring at RT for 60min. After centrifugation (20000g, 45min, 5°C), unwanted proteins were removed from the supernatant by 30% ammonium sulfate precipitation. The supernatant was further purified using hydrophobic interaction chromatography (Phenylsepharose 6, fast flow, GE Healthcare) using 25mM imidazole, 1.25M ammonium sulfate, pH 7.5 as binding buffer and 25mM imidazole pH7.4 for the elution gradient. Fractions were analyzed on a 15% acrylamide gel by SDS PAGE and Phl p 6 containing fractions were pooled and dialyzed three times against 25mM imidazole, 4% 2-propanol, pH 7.4. Dialyzed pooled fractions were applied to anion exchange chromatography (DEAE sepharose fast flow, GE Healthcare) using 20mM imidazole, 4% 2-propanol, pH7.4 for equilibration and washing. Elution was performed with a NaCl gradient in the same buffer. Fractions containing Phl p 6 were finally purified by size exclusion chromatography using a 100/60 sephacryl S300HR (GE Healthcare) column equilibrated with 2mM sodium phosphate buffer, pH 7.4.

^15^N and ^15^N/^13^C labeled protein samples for NMR spectroscopy were recombinantly produced using M9 minimal media supplemented with ^15^NH_4_Cl and ^13^C-D-glucose and purified as described above.

#### Molecular Dynamics and Protein flexibility

We performed molecular dynamics simulations of wild-type Phl p 6 and the four *in silico* selected mutants to profile the respective conformational ensembles and intrinsic flexibilities. In order to achieve broad conformational sampling, we used accelerated molecular dynamics simulations (aMD). For the wild-type, the starting structure for the simulation was obtained from the available crystal structure 1NLX. We note that in this structure, only 104 of the total 111 residues of the allergen are resolved. In detail, four C-terminal residues as well as three N-terminal residues (including the starting methionine) are not resolved. For the mutants, starting structures were modelled based on the wild-type structure. The respective point mutations were introduced with the program MOE (molecular operating environment) ^5^, followed by local minimization. Structures were protonated at pH 7.0 with the tool protonate3d as implemented in MOE. Topology and starting coordinate files were generated with the LEaP module of AmberTools 17 ^6^ using the ff99SB-ILDN force field ^7^. Each protein was placed in a truncated octahedral TIP3P water box^8^ with a minimum wall distance of 10 Å. After an elaborate equilibration protocol, consisting of subsequent relaxing, heating and cooling steps^9^short conventional molecular dynamics simulations of 100 ns length were performed to obtain acceleration parameters ^10^. Subsequent accelerated MD simulations were performed for 1000 ns per system. Simulations were carried out in the NpT ensemble, using a Langevin thermostat t^11^ with a collision frequency of 2 ps^-1^ to keep a constant temperature of 310 K, as well as a Berendsen barostat^12^ with a relaxation time of 2 ps to keep the system at atmospheric pressure. Bonds involving hydrogen bonds were constrained with the SHAKE algorithm to allow the use of 2 fs time-step ^13^. A van der Waals cutoff of 10 Å was used and long-range electrostatics were treaded the particle-mesh Ewald method (PME)^14^. All simulations were carried out with the GPU acceleration of the pmemd module of AMBER 17 on our in-house cluster. Snapshots were saved each 10 ps.

All trajectories were reweighted with a McLaurin series to the 10^th^ order prior to analysis ^10, 15^. Analyses were carried out with the program cpptraj ^16^ of AmberTools 17 and in-house python scripts. Local flexibilities were characterized by calculating the dihedral entropies resulting from backbone torsion probability profiles ^17^. Structural visualizations were achieved by calculating the residues-wise differences in dihedral entropy for each system, using the wild-type as a reference and mapping the respective differences onto the respective crystal structures using the program PyMol ^18^.

#### Protein folding and stability

Protein folding in solution was monitored by circular dichroism spectroscopy using a JASCO J-815 spectropolarimeter equipped with a PTC-423S Peltier-type single position cell holder (Jasco). Proteins were dissolved to 10 µM in 2 mM sodium phosphate buffer pH 7.4 and spectra were recorded from 190 to 260 nm. Thermal denaturation was monitored at 222 nm from 20 to 90°C with a temperature slope of 1°C/min.

All NMR measurements were carried out with 0.75 mM Phl p 6 in 10 mM sodium phosphate buffer pH 7.0 with 10% D_2_O. For protein assignment, standard triple-resonance methodology was employed. ^15^N HSQC spectra ^19^ of wild-type Phl p 6  and L89G and E39L mutants were measured in a temperature range between 20°C and 70°C in steps of 5°C. Chemical shift changes due to the change in temperature were recorded using 500 µM sodium trimethylsilylpropanesulfonate (TMS) as reference substance (0 ppm at all temperatures).

#### Protein crystallization, data collection and structure determination

For initial crystallization screening the sitting-drop vapor-diffusion method was applied, utilizing a Hydra II Plus One (Matrix) liquid-handling system. 0.2 µl Phl p 6 mutant proteins at a concentration of 10 mg ml^−1^ were mixed with 0.2 µl screen solution from different commercial screens and equilibrated against 60 µl reservoir solution in a 96-well-plate (Art Robbins Instruments) at 293 K. Crystals were observed in condition consisting of 0.05 M zinc acetate dehydrate, 20% (w/v) polyethylene glycol (PEG) 3350 for Phl p 6 S46Y. Fine screening was carried out using different protein to drop ratio 1:1, 1:1.5, 1:2, PEG3350 concentration 10-25% in hanging drop at 293K. Phl p 6 S46Y mutant crystals were observed in condition consisting of 0.05 M zinc acetate dehydrate, 10% (w/v) PEG3350 with a protein to drop ratio of 1:1. Micro-seeding using Phl p 6 S46Y crystals was performed to ease the crystallization of other mutants ^20^. The crystals were cryo-protected with 10% glycerol and flash frozen in liquid nitrogen. Diffraction datasets were collected at the ESRF beamline ID29 and processed using iMosflm ^21^ and Scala in the CCP4 software suite ^22^. The structure was solved by molecular replacement using the existing Phl p 6 structure (PDB: 1NLX). Molecular replacement was done using CCP4 software suite. Structure models were built using COOT ^23^ and refined using Phenix ^24^. Anomalous data sets were collected at the Zinc K-absorption edge and data processing was performed as described above. The two data sets were then combined and scaled using Scaleit in the CCP4 software suite. The anomalous difference was calculated using Sftools in the CCP4 software suite. Finally, the anomalous difference Fourier maps were calculated with FFT in the CCP4 software suite. The coordinates of Phl p 6 S46Y have been deposited in the Protein Data Bank under the entry code 6TRK, see supplementary Table 2 for crystallographic data and refinement statistics.

#### Endolysosomal degradation – peptide analysis

Intact proteins as well as peptides generated from Phl p 6 and the mutants were assessed by high-performance liquid chromatography-mass spectrometry (HPLC-MS) using a Q Exactive™ Hybrid Quadrupole-Orbitrap™ mass spectrometer (Thermo Fisher Scientific) online hyphenated to an UltiMate™ RSLC nano system (Thermo Fisher Scientific) by means of a nano-electrospray ionization source. Prior to analysis, samples were purified using Pierce™ C18 tips (Thermo Fisher Scientific) as described in the manufacturer’s manual.

For HPLC-MS analysis of intact proteins, 250 ng were injected onto a Waters X Bridge Protein BEH C4 2.1x150mm column with 3.5 μm particles and a pore size of 300Å (Waters, Milford, MA, USA). Water (A) and acetonitrile (B; Sigma Aldrich) each containing 0.10% (v/v) formic acid (FA) were used as eluents. At a flow rate of 200 µL/min proteins were eluted by applying a linear gradient of 20.0–60.0% B over 3 min. The column temperature was set to 60°C. The mass spectrometer was operated in positive ionization mode. Full scans were performed at a scan range of m/z 500 to 1,500 at a resolution of 140,000 (at m/z = 200). The AGC-target was set to 1 × 10^6^ with a maximum injection time (IT) of 100 ms.

To identify peptides resulting from endolysosomal degradation, 600 ng were injected onto a self-packed 200 x 0.1 mm Hypersil GOLD™ aQ C18 (Thermo Fisher Scientific) capillary column with 3.0 µm particles. Peptides were separated at a flow rate of 350 nl/min using a linear gradient of 5.0-40.0% B in 60 min at 50°C. For MS detection, survey scans were performed for a scan range of m/z 370-2,000 at a resolution of 70,000 (at m/z = 200). The AGC target was set to 1 x 10^6^ with a maximum IT of 120 ms. The 15 most intense ions were selected for fragmentation using a normalized collision energy of 29. Data dependent MS2 scans were recorded at a resolution of 35,000 with an AGC target of 2 x 10^5^ and a maximum IT of 100 ms.

For data analysis Xcalibur 3.0 (Thermo Fisher Scientific), Proteome Discoverer 1.4 (Thermo Fisher Scientific) and GPMAW (Lighthouse data, Denmark) were used.

#### Generation of Phl p 6–specific T cell hybridomas

Splenocytes from Phl p 6 immunized mice were cultured in Opti-MEM + GlutaMAX (Gibco), 5% FCS, 100U/mL penicillin, 100µg/mL streptomycin, with the immundomaint peptide 31 (AA 92-106) at a final concentration of 3µg/mL. One week later, dead cells and peptides were removed by ficoll density gradient centrifugation and cells were incubated with 20U/ml recombinant human IL-2 for another week. After removal of IL-2 by washing, T cells were co-cultured with syngenic irradiated feeder cells (splenocytes from naïve BALB/c mice) with 6µg/mL of the respective peptide for 8 days. On day 8, T cells were harvested and fused with mouse thymus lymphoma cell line BW5147.G.1.4 (ATCC) according to standard protocols for generation of T cell hybridomas.^25^

#### Generation of BMDCs

Bone marrow was harvested from femur and tibia of BALB/c mice. For generation of GM-CSF BMDCs, 10 mL of a 2x10^5^ cells/mL bone marrow cell suspension were plated into non-cell culture treated petri dishes and incubated in the presence of murine GM-CSF (Immunotools, Germany) for 6-8 days. On day 3, 10 mL of GM-CSF containing culture medium (RPMI1640 supplemented with 20 ng/mL GM-CSF, 100 U/mL penicillin, 100 µg/mL streptomycin, 2 mM L-Glutamine, 10% FCS and 50 µM 2-mercaptoethanol) was added to each well, and on day 5, 10 mL medium was replaced.

#### Lymphocyte cultures

All mice received terminal anesthesia (100 mg/kg Ketamine + 3 mg/kg Xylazine + 3 mg/kg Acepromazine) by i.p. injection, blood was sampled from retroorbital sinus, and mice were sacrificed by cervical dislocation. Spleens were aseptically removed and transferred into 40mm Petri dishes containing 500µL DPBS. Spleens were homogenized using the back of a sterile plunger from a 2mL syringe and the suspensions were transferred into 1.5mL tubes. After 3-5 min incubation at room temperature (until debris had settled), the monodisperse cell suspension was transferred into a 15mL tube, pre-filled with 7mL of ACK red blood cell lysis buffer (0.15M NH_4_Cl, 10mM KHCO_3_, 0.1mM NA_2_EDTA, pH 7.2-7.4) and incubated for 7min at RT. Tubes were filled up with 6mL of DPBS, centrifuged for 5min at 260g at RT, and the pellets were washed with 5mL of DPBS. After centrifugation, the pellets were resuspended in 10mL warm DPBS containing 1µM eFluor 450 cell proliferation dye (eBioscience/Thermo Fisher Scientific). After incubation for 10 min at 37°C, labeling was stopped by addition of 10% FCS. After two washing steps with DPBS, lymphocytes were cultured in T-cell medium medium (RPMI, 10% FCS, 25mM HEPES, 2mM L-Glu, 100µg/mL streptomycin, 100U/mL penicillin) and counted using a CEDEX XS Cell Analyzer (Roche). 3x10^5^ cells per well were incubated in U-bottom tissue culture plates with 20µg/mL Phl p 6 or the mutant proteins or 10µg/mL of individual peptides from a 15mer library (GenScript, NJ, USA) with an offset of 3 amino acids in a total volume of 150µL per well. After 4 days, splenocyte supernatants were removed and cells were harvested for flow cytometry analysis.

#### Epitope mapping

Restimulated splenocytes were transferred to V-bottom plates and stained in DPBS for 10 min at RT with fixable viability dye eFluor 780 (1:5000, eBioscience) and anti-mouse CD4 PerCp/Cy5.5 (1:200, BioLegend, SanDiego, USA, clone GK1.5) and finally analyzed on a Cytoflex S flow cytometer (Beckmann Coulter). Proliferating cells among live CD4+ T cells were identified by their reduced fluorescence on the eFluor 450 channel. Peptides were considered immunoreactive if the stimulation index (% proliferating cells stimulated with peptide / % proliferating cells without stimulation) was greater than two.

#### Measurement of Phl p 6-specific serum immunoglobulins and cell bound IgE (RBL assay and BAT)

A luminometric ELISA was performed to determine the production of Phl p 6-specific serum IgG1 and IgG2a. 96-well plates (flat white chimney; Greiner) were coated with 1 µg/mL recombinant Phl p 6 or the mutant proteins diluted in PBS (50 µL per well) overnight at 4°C. The coating solution was discarded and 200 µL of blocking buffer (2% skim milk, blotting grade, and 0.1% Tween®-20 in PBS) were added to each well and incubated 1 h at RT. The plate was washed with PBS-T (PBS with 0.1% Tween-20) using an automated plate washer (Tecan 96PW Microplate Washer). Meanwhile, serum dilutions (1:100 for IgG2a and 1:100,000 for IgG1) were prepared in blocking buffer. 50 µL of each serum dilution was added to the wells and incubated for 2 h at 37°C in a humid chamber. After another washing step, 50 µL of the secondary antibody (goat anti-mouse IgG1-HRP diluted 1:1000, Bio-Rad; rabbit anti-mouse IgG2a-HRP diluted 1:2000, antibodies-online.com) were added in blocking buffer. The plate was incubated for 1 h at RT. After a final washing step, 50 µL of an ELISA BM chemiluminescence substrate (Roche) was added and after 2-3 min the luminescence was measured in relative light units (RLU) using a plate reader (Tecan inifinite 200Pro; integration time of 1000 ms and attenuation set to automatic).

As a correlate for biologically active, specific IgE in sera, a β-hexosaminidase release assay was performed. Briefly, rat basophil leukemia (RBL-2H3) cells were seeded in 96-well tissue culture plates and incubated overnight at 37°C, 5% CO_2,_ 95% humidity. The next day, sera were added for passive sensitization at indicated dilutions. Background wells and wells for maximum release remained untreated. After 2h of incubation, cells were washed three times with Tyrode’s buffer containing 1 % BSA and incubated for 1h with 10ng/mL of Phl p 6 or the mutant proteins in Tyrode’s buffer for stimulation. 10% Triton X-100 was added to some wells for maximum release. Supernatants containing released mediators including β-hexosaminidase were transferred to fresh plates and assay solution (4-MUG diluted in 0.1M citrate buffer, pH 4.5) was added for 1h. The reaction was stopped by addition of 100µL glycine buffer (0.2M glycine, 0.2M NaCl, pH 10.7). Fluorescence intensity was measured on a TECAN plate reader at λ_ex_:360nm and λ_em_:465nm. Percentage of specific lysis was calculated by the formula (rfu – rfu_back_)/(rfu_max_ – rfu_back_)*100.

Cell bound IgE was measured *ex vivo* using a basophil activation test. On the day of sacrifice, blood was taken from the retro orbital sinus and mixed with 1/10 vol. of Li-Heparin (10mg/mL in DPBS) to prevent coagulation. 30µL of these blood samples were mixed with 30µL of RPMI with an equimolar mix of Phl p 6 WT, L89G, and S46Y and incubated for 2h at 37°C, 5% CO_2_, 95% humidity. Final concentration of Phl p 6 in stimulated wells was 10ng/mL. After 2h, stimulation was stopped by putting the plates on ice, and all subsequent steps were performed with ice-cold solutions and samples were kept on ice. Samples were washed with 120µL FACS buffer (PBS, 1% BSA, 2mM EDTA) and cell pellets (5min centrifugation at 260 g) were resuspended in 30µL of FACS buffer containing anti-IgE-FITC (BioLegend, clone RME-1, 1:200), anti-CD4-PerCp-Cy5.5 (BioLegend, clone GK1.5, 1:200), anti-CD19-PE/Cy7 (BioLegend, clone 6D5, 1:200), and anti-CD200R-APC (eBioscience/Thermo Fisher Scientific, clone OX110, 1:200). Samples were stained for 20min on ice and washed with 100µL FACS buffer. After 5 min centrifugation at 260g, pellets were resuspended in 150µL red blood cell lysis buffer (eBioscience/Thermo Fisher Scientific) and incubated for 5min at RT. Cells were centrifuged for 5min at 400g and pellets were washed two times with FACS buffer. Finally, cell pellets were resuspended in 50µL FACS buffer and analyzed on a FACS Canto II flow cytometer. Basophils were gated as IgE^high^ CD19^neg^ CD4^neg^ cells and activation status was assessed by the median fluorescence intensity of CD200R as previously described ^26^.

### Supplementary Results:

#### Identification of dominant T cell epitopes

The dominant T cell epitopes of Phl p 6 were identified by culturing splenocytes from mice immunized with Phl p 6 together with 33 overlapping peptides covering its entire sequence (15mers, 3AA overlap). By determining the proliferation rate of CD4+ T cells, two major (peptides 22 and 31) and one minor (peptide 28) T cell epitopes were identified (suppl. Fig. 1). Sequences of the respective T cell epitopes are shown in supplementary Table 1.

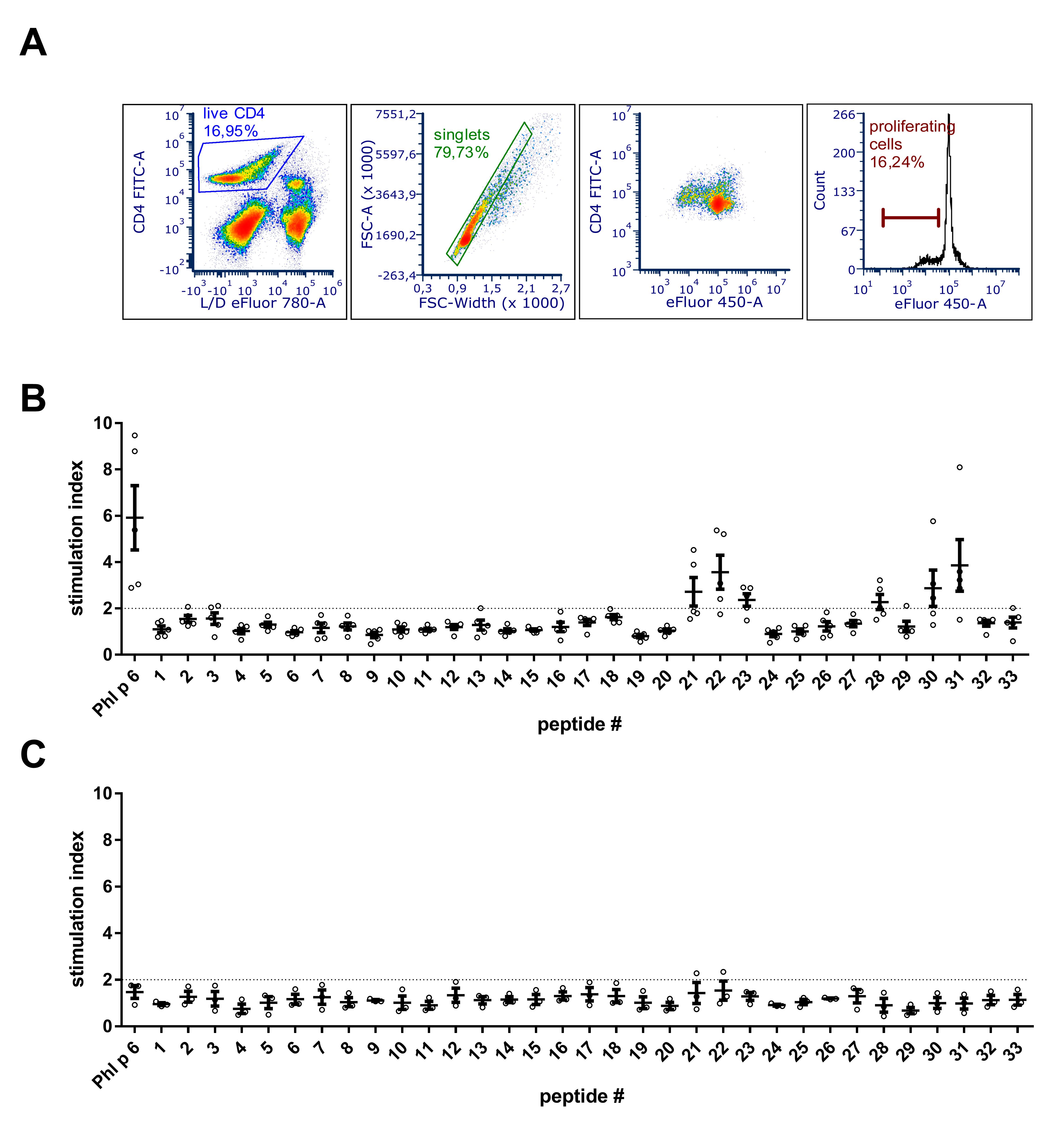

Supplementary Figure 1. Epitope mapping. Splenocytes from Phl p 6 immunized BALB/c mice (B) or naïve control mice (C) were restimulated with Phl p 6 or 33 individual peptides covering its entire sequence. Proliferation of splenocytes was determined by eFluor 450 proliferation assay and data are shown as stimulation index (%proliferation peptide / %proliferation medium). A stimulation index above 2 (dotted line) was considered to represent a significant signal. Data are shown for individual mice (n=5) and means±SEM. (A) Gating strategy: live CD4 cells were gated followed by exclusion of doublets. Proliferating T cells were identified by their reduced levels of eFluor 450 dye.

Supplementary Table 1 – BALB/c T cell epitopes of Phl p 6

| Peptide # | Sequence | AA position |
| --- | --- | --- |
| 22 | DEVYNAAYNAADHAA | 65-79 |
| 28 | KYEAFVLHFSEALRI | 82-96 |
| 31 | SEALRIIAGTPEVHA | 92-106 |

#### Characterization of recombinant Phl p 6 and its mutants

Purified recombinant Phl p 6 and its mutants were analyzed on a 15% acrylamide gel by SDS-PAGE and Coomassie staining (suppl. Fig. 2A). To confirm the amino acid substitution the exact mass of the mutant proteins was verified by MS. The obtained spectra were deconvoluted using the Xtract function of Xcalibur 3.0 software (Thermo Fisher Scientific) and the experimentally determined mass was compared to the theoretical mass of the WT and the mutants (suppl. Fig. 2B).

CD spectra recorded from 190 to 260 nm confirmed that the overall three dimensional structure was not altered by introduction of the stabilizing or destabilizing mutations (suppl. Fig. 2C).

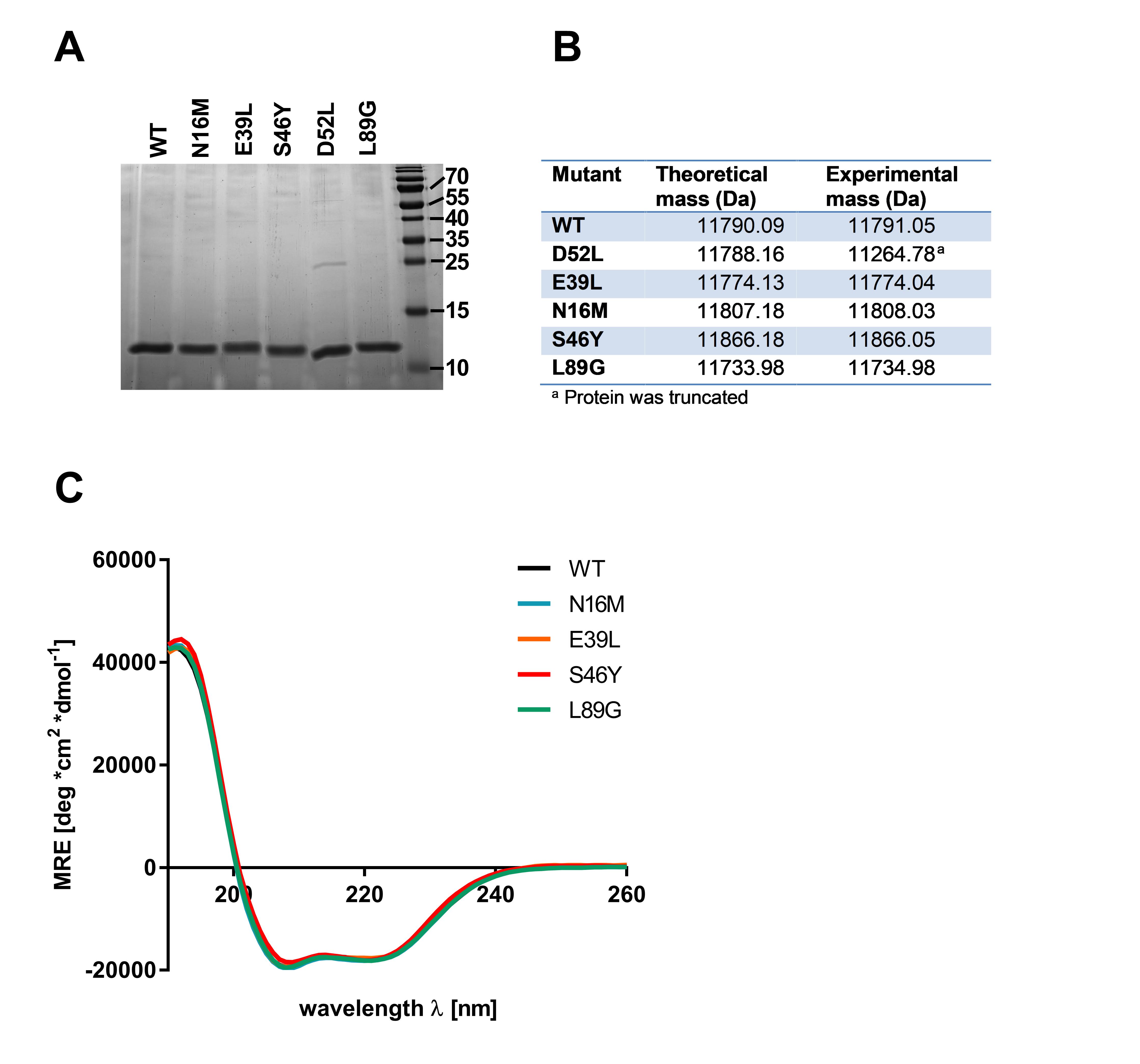

Supplementary Figure 2. Characterization of recombinant Phl p 6 (WT) and its mutants (D52L, E39L, N16M, S46Y, L89G). A) SDS-PAGE B) Theoretical and experimental average mass as determined by HPLC-MS C) Full CD spectra measured at room temperature, pH 7.4 and mean residue ellipticity (MRE) is shown.

**Supplementary Table 2.** **Crystallographic data and refinement statistics.**

| Structure | S46Y | S46Y, Zn peak | S46Y, Zn remote |
| --- | --- | --- | --- |
| Data collection  Wavelength (Å)  Unit cell parameters  *a,b,c (Å)*  *α,β,γ (degrees)*  Space group  Solvent content (%)  Protein chains in AU  Resolution range (Å)  Highest resolution shell (Å)  Unique reflections  Redundancy  Completeness (%)  **R*_merge_  *R*_meas_  Average *I/σ(I)* | 0.9763  47.83, 73.51, 76.12  *α=β=γ=*90  P22_1_2_1_  56.64  2  39.94-1.60  1.69-1.60  34555 (5013)  3.5 (3.1)  96.9 (97.5)  0.070 (0.439)  0.082 (0.523)  9.3 (2.3) | 1.2815  47.49, 73.13, 75.78  *α=β=γ=*90  P22_1_2_1_  56.64  2  47.79-1.61  1.69-1.61  35203 (5075)  7.2 (7.3)  99.9 (100)  0.083 (0.794)  0.104 (0.926)  10.9 (2.2) | 1.2848  47.65, 73.29, 75.91  *α=β=γ=*90  P22_1_2_1_  56.64  2  47.65-1.61  1.70-1.61  35198 (5072)  7.3 (7.4)  99.9 (100)  0.064 (0.424)  0.074 (0.493)  14.3 (3.7) |
| Refinement  *R*_work_ (%)  *R*_free_ (%)  Mean *B* value (Å^2^)  B from Wilson plot (Å^2^)  RMSD bond length (Å)  RMSD bond angles (°)  No. of amino acid residues  No. of water molecules  No. of metal ions | 15.52  18.27  29  16.9  0.007  0.859  108  303  4 |  |  |
| Ramachandran plot  Most favored regions (%)  Allowed regions (%) | 99.55  0.45 |  |  |

Values of the highest resolution shell are given in parentheses.

*R_merge_ = Σ_h_Σ_l_ |*I*_hl_ − ⟨*I*_h_⟩|/Σ_h_Σ_l_ |⟨*I*_h_⟩|

#### Molecular Dynamics

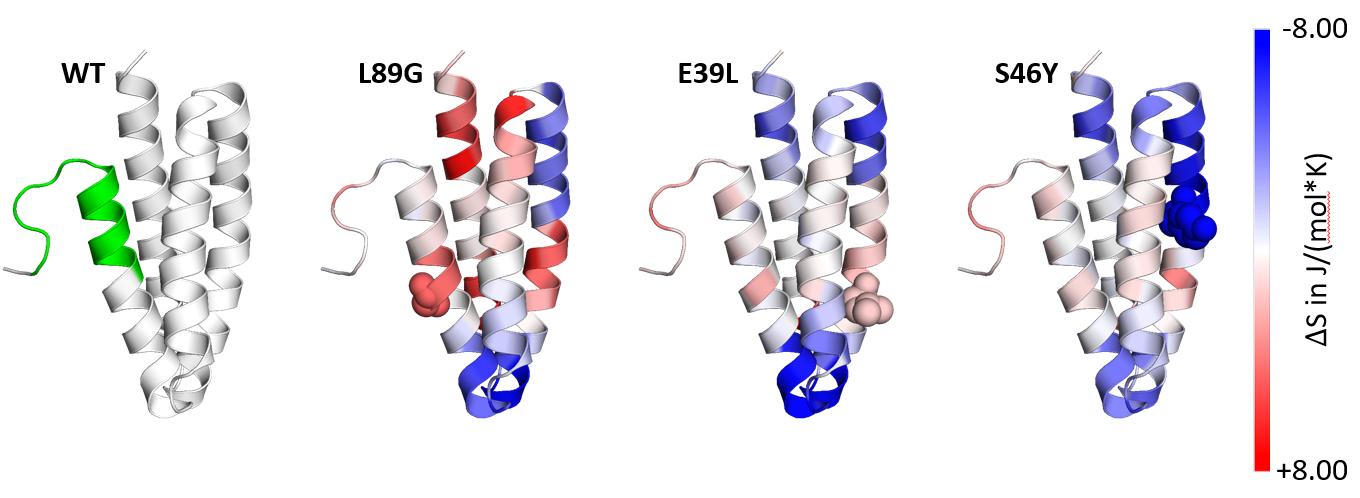

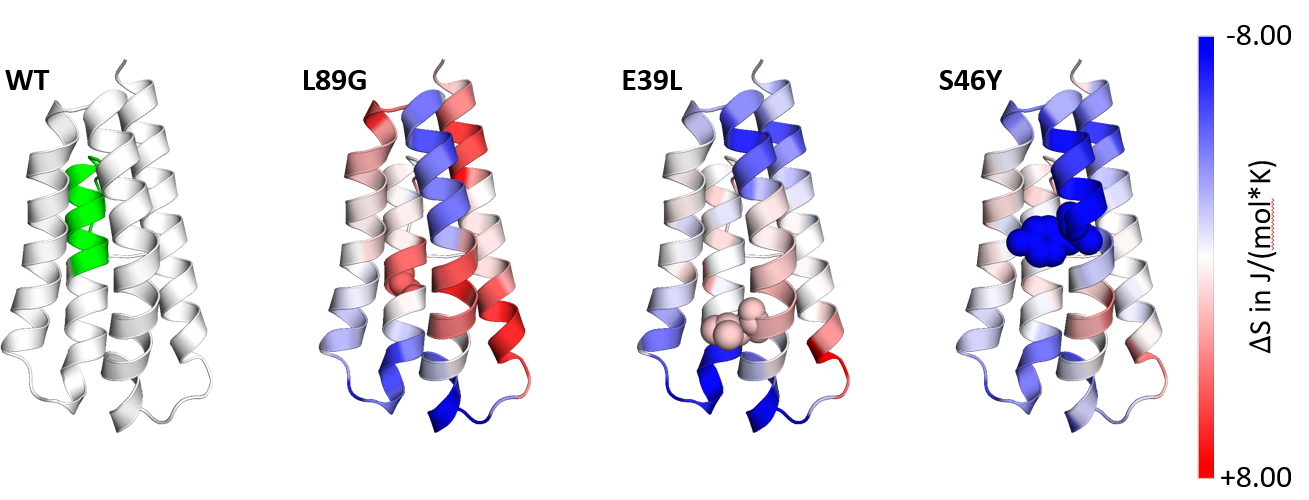

Supplementary Figure 3. Dihedral Entropies calculated from aMD simulations to predict localized flexibility. The immunodominant epitope AA92-106 is shown in green. Red indicates increased flexibility compared to the WT molecule and blue indicates reduced flexibility.

#### Degradome Assay

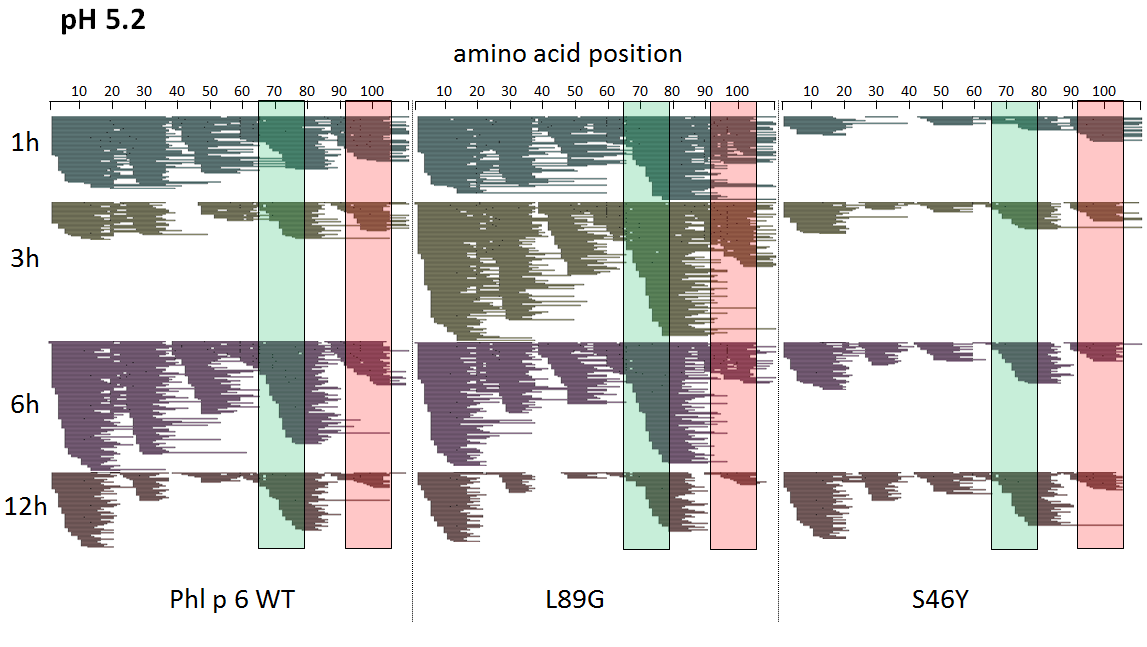

Supplementary Figure 4. Degradation pattern of Phl p 6 wild type (WT) and the mutants L89G and S46Y after incubation with an endolysosomal extract at pH 5.2. Peptideswere identified by LC-MS/MS and are displayed color coded for different incubation times (1, 3, 6, and 12 hours). Amino acid regions corresponding to the immunodominant epitopes 65-79 and 92-106 of Phl p 6 are highlighted in green and red, respectively.

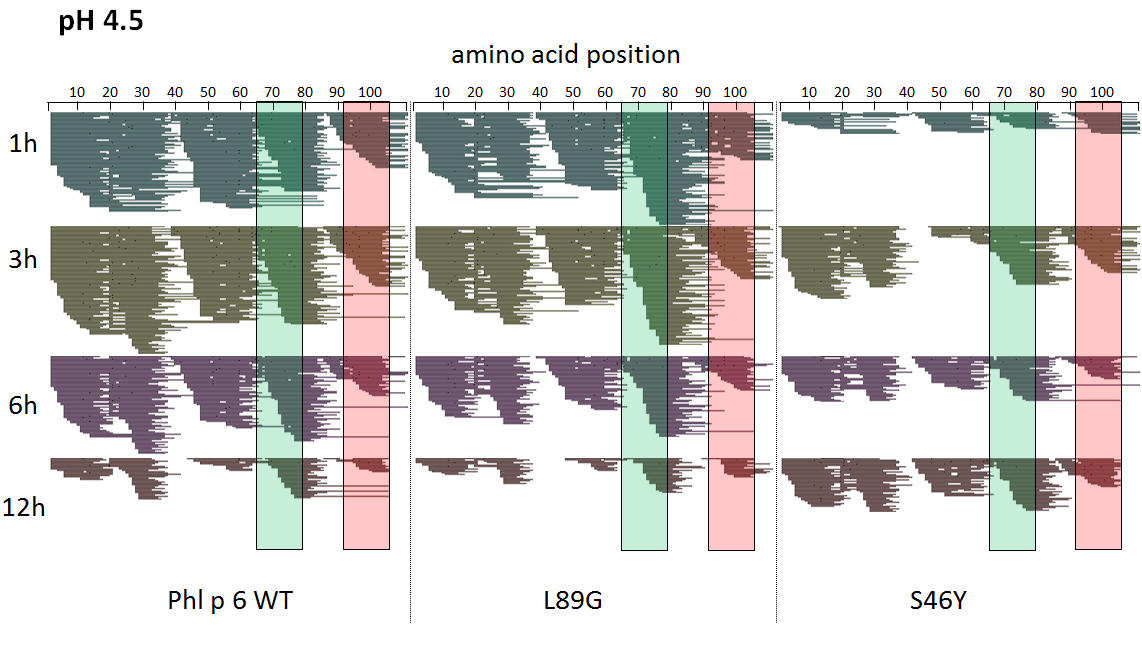

Supplementary Figure 5. Degradation pattern of Phl p 6 wild type (WT) and the mutants L89G and S46Y after incubation with an endolysosomal extract at pH 4.5. Peptides were identified by LC-MS/MS and are displayed color coded for different incubation times (1, 3, 6, and 12 hours). Amino acid regions corresponding to the immunodominant epitopes 65-79 and 92-106 of Phl p 6 are highlighted in green and red, respectively.

#### Cross reactivity of induced antibodies

To test whether stabilizing or destabilizing mutations changed the fine specificity of Phl p 6 paratopes, sera from mice immunized with Phl p 6 WT, S46Y, and L89G were tested against the different proteins by ELISA. As shown in suppl. Fig. 6A and B, antibodies raised against the WT protein showed significantly lower reactivity against the destabilized mutant L89G as well as the stabilized mutant S46Y. Sera from mice immunized with the stabilized mutant S46Y showed much lower IgG1 reactivity against the destabilized mutant L89G, and only slightly lower reactivity against the WT protein. The differences in binding of IgG2a to the different proteins were much lower and mostly not significant (suppl. Fig. 6B).

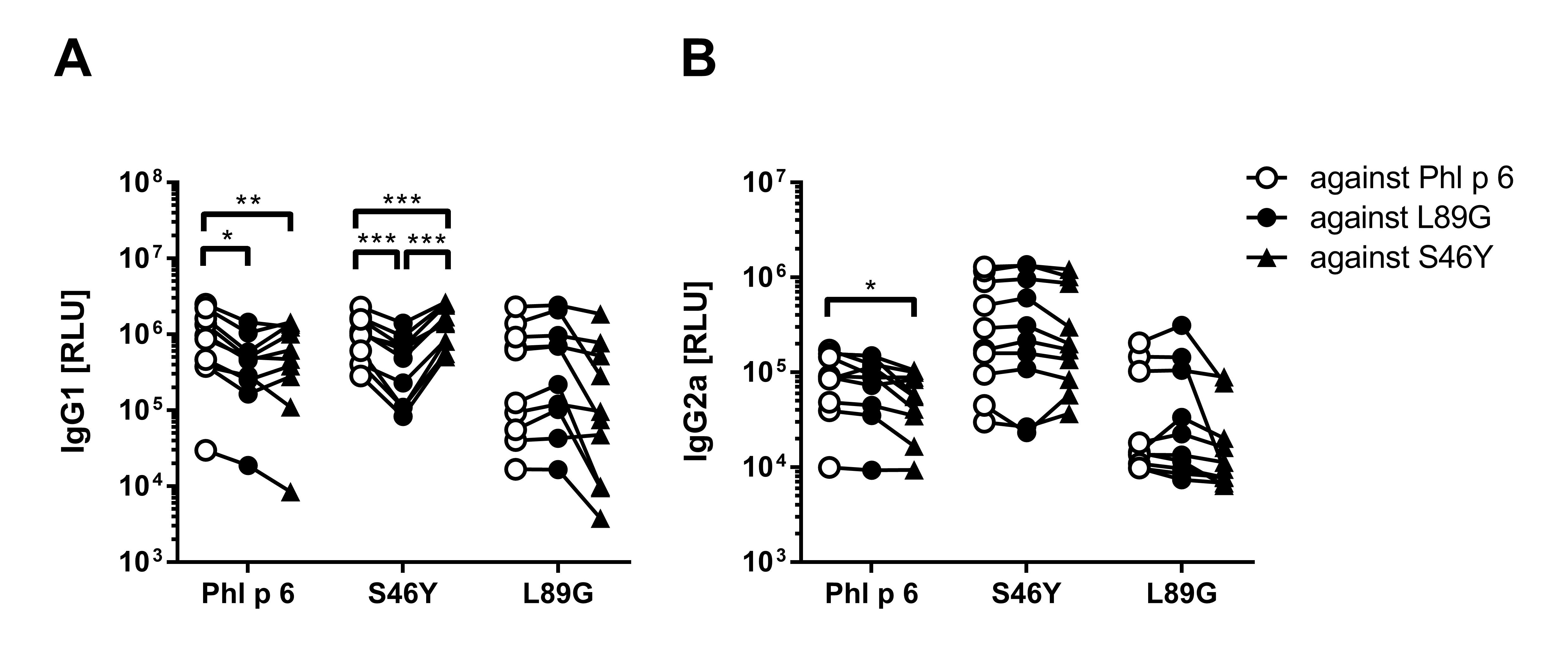

Supplementary Figure 6. Sera from mice immunized with Phl p 6, L89G, and S46Y were analyzed for their reactivity against Phl p 6 WT and the different mutant proteins. Data are shown as relative light units (RLU) of a luminometric IgG1 (A) or IgG2a (B) ELISA. Statistical differences in binding were assessed by two-way RM ANOVA followed by Tukey’s post hoc test. * P<0.05, ** P<0.01, *** P<0.001.

### References

1.	Laimer J, Hofer H, Fritz M, Wegenkittl S, Lackner P. MAESTRO--multi agent stability prediction upon point mutations. BMC Bioinformatics 2015; 16:116.

2.	Pettersen EF, Goddard TD, Huang CC, Couch GS, Greenblatt DM, Meng EC, et al. UCSF Chimera--a visualization system for exploratory research and analysis. J Comput Chem 2004; 25:1605-12.

3.	Webb B, Sali A. Comparative Protein Structure Modeling Using MODELLER. Curr Protoc Protein Sci 2016; 86:2 9 1-2 9 37.

4.	Studier FW. Protein production by auto-induction in high density shaking cultures. Protein Expr Purif 2005; 41:207-34.

5.	Molecular Operating Environment (MOE). 1010 Sherbooke St. West, Suite #910, Monteal, QC, Canada, H3A 2R7: Chemical Computing Group, 2017.

6.	Case DA, Cerutti DS, Cheatham III TE, Darden TA, Duke RE, Giese TJ, et al. AMBER 2017; 2017. University of California, San Francisco.

7.	Lindorff-Larsen K, Piana S, Palmo K, Maragakis P, Klepeis JL, Dror RO, et al. Improved side-chain torsion potentials for the Amber ff99SB protein force field. Proteins: Structure, Function, and Bioinformatics 2010; 78:1950-8.

8.	Jorgensen WL, Chandrasekhar J, Madura JD, Impey RW, Klein ML. Comparison of simple potential functions for simulating liquid water. The Journal of Chemical Physics 1983; 79:926-35.

9.	Wallnoefer HG, Handschuh S, Liedl KR, Fox T. Stabilizing of a Globular Protein by a Highly Complex Water Network: A Molecular Dynamics Simulation Study on Factor Xa. The Journal of Physical Chemistry B 2010; 114:7405-12.

10.	Pierce LCT, Salomon-Ferrer R, Augusto F. de Oliveira C, McCammon JA, Walker RC. Routine Access to Millisecond Time Scale Events with Accelerated Molecular Dynamics. Journal of Chemical Theory and Computation 2012; 8:2997-3002.

11.	Adelman SA, Doll JD. Generalized Langevin equation approach for atom/solid‐surface scattering: General formulation for classical scattering off harmonic solids. The Journal of Chemical Physics 1976; 64:2375-88.

12.	Berendsen HJC, Postma JPM, Gunsteren WFv, DiNola A, Haak JR. Molecular dynamics with coupling to an external bath. The Journal of Chemical Physics 1984; 81:3684-90.

13.	Ryckaert J-P, Ciccotti G, Berendsen HJC. Numerical integration of the cartesian equations of motion of a system with constraints: molecular dynamics of n-alkanes. Journal of Computational Physics 1977; 23:327-41.

14.	Darden T, York D, Pedersen L. Particle mesh Ewald: An N⋅log(N) method for Ewald sums in large systems. The Journal of Chemical Physics 1993; 98:10089-92.

15.	Miao Y, Sinko W, Pierce L, Bucher D, Walker RC, McCammon JA. Improved Reweighting of Accelerated Molecular Dynamics Simulations for Free Energy Calculation. Journal of Chemical Theory and Computation 2014; 10:2677-89.

16.	Roe DR, Cheatham TE. PTRAJ and CPPTRAJ: Software for Processing and Analysis of Molecular Dynamics Trajectory Data. Journal of Chemical Theory and Computation 2013; 9:3084-95.

17.	Kamenik AS, Kahler U, Fuchs JE, Liedl KR. Localization of Millisecond Dynamics: Dihedral Entropy from Accelerated MD. Journal of Chemical Theory and Computation 2016; 12:3449-55.

18.	Schrodinger, LLC. The PyMOL Molecular Graphics System, Version 1.8.6.0. 2017.

19.	Kay LE, Keifer P, Saarinen T. Pure Absorption Gradient Enhanced Heteronuclear Single Quantum Correlation Spectroscopy with Improved Sensitivity. Journal of the American Chemical Society 1992; 114:10663-5.

20.	Stewart PDS, Kolek SA, Briggs RA, Chayen NE, Baldock PFM. Random Microseeding: A Theoretical and Practical Exploration of Seed Stability and Seeding Techniques for Successful Protein Crystallization. Crystal Growth & Design 2011; 11:3432-41.

21.	Powell HR, Battye TGG, Kontogiannis L, Johnson O, Leslie AGW. Integrating macromolecular X-ray diffraction data with the graphical user interface iMosflm. Nat Protoc 2017; 12:1310-25.

22.	Winn MD, Ballard CC, Cowtan KD, Dodson EJ, Emsley P, Evans PR, et al. Overview of the CCP4 suite and current developments. Acta Crystallogr D Biol Crystallogr 2011; 67:235-42.

23.	Emsley P, Lohkamp B, Scott WG, Cowtan K. Features and development of Coot. Acta Crystallogr D Biol Crystallogr 2010; 66:486-501.

24.	Adams PD, Afonine PV, Bunkoczi G, Chen VB, Davis IW, Echols N, et al. PHENIX: a comprehensive Python-based system for macromolecular structure solution. Acta Crystallogr D Biol Crystallogr 2010; 66:213-21.

25.	Kruisbeek AM. Production of mouse T cell hybridomas. Curr Protoc Immunol 2001; Chapter 3:Unit 3.14.

26.	Korotchenko E, Moya R, Scheiblhofer S, Joubert IA, Horejs-Hoeck J, Hauser M, et al. Laser facilitated epicutaneous immunotherapy with depigmented house dust mite extract alleviates allergic responses in a mouse model of allergic lung inflammation. Allergy 2019.
